# Genetics of skeletal proportions in two different populations

**DOI:** 10.1101/2023.05.22.541772

**Authors:** Eric Bartell, Kuang Lin, Kristin Tsuo, Wei Gan, Sailaja Vedantam, Joanne B. Cole, John M Baronas, Loic Yengo, Eirini Marouli, Tiffany Amariuta, Zhengming Chen, Liming Li, GIANT consortium, China Kadoorie Biobank Collaborative Group, Nora E Renthal, Christina M. Jacobsen, Rany M Salem, Robin G Walters, Joel N Hirschhorn

## Abstract

Human height can be divided into sitting height and leg length, reflecting growth of different parts of the skeleton whose relative proportions are captured by the ratio of sitting to total height (as sitting height ratio, SHR). Height is a highly heritable trait, and its genetic basis has been well-studied. However, the genetic determinants of skeletal proportion are much less well-characterized. Expanding substantially on past work, we performed a genome-wide association study (GWAS) of SHR in ∼450,000 individuals with European ancestry and ∼100,000 individuals with East Asian ancestry from the UK and China Kadoorie Biobanks. We identified 565 loci independently associated with SHR, including all genomic regions implicated in prior GWAS in these ancestries. While SHR loci largely overlap height-associated loci (P < 0.001), the fine-mapped SHR signals were often distinct from height. We additionally used fine-mapped signals to identify 36 credible sets with heterogeneous effects across ancestries. Lastly, we used SHR, sitting height, and leg length to identify genetic variation acting on specific body regions rather than on overall human height.

## Introduction

Human height has long been studied as a model polygenic phenotype due to its high heritability (Yang et al. 2015; Lango Allen et al. 2010; Wood et al. 2014; Loic Yengo et al. 2018; Loïc Yengo et al. 2022), polygenicity (O’Connor et al. 2019), and ease of accurate measurement. Recent work from the GIANT consortium, combining data from 5.4 million individuals across 5 major population groups, maps genomic regions that account for nearly all of the heritability attributable to common variation (h^2^_snp_, estimated to be 0.514), identifying over 12,000 signals associated with height (Loïc Yengo et al. 2022).

Standing height reflects the sum of the sizes of different skeletal components, whose relative proportions vary across individuals and populations. Despite the genetics of height being well-studied, much less is known about the genetics of body proportion (Chan et al. 2015; Kun et al. 2023). Here, we hypothesize that investigating genetics of body proportion will provide insights into our understanding of human skeletal growth. Furthermore, as height is frequently used as a model polygenic trait, the availability of traits that are components of height offers a setting for using genetics to study a set of polygenic traits with partially overlapping underlying biology.

To better understand the genetics and biology of height, multiple related phenotypes could potentially provide additional insights. We have previously studied sitting height ratio (SHR), the ratio of sitting height to standing height (Chan et al. 2015), as well as protein levels for growth related genes to begin to understand the underlying biology at selected height loci (Bartell et al. 2020). Our prior GWAS of SHR (Chan et al. 2015), in 3,545 African American individuals and 21,590 individuals of European ancestry, identified 6 genome-wide significant associations, including at the locus encoding the growth-related protein IGFBP-3. The substantial heritability of SHR (estimated as 26% previously, (Chan et al. 2015)) suggests that a larger sample size, combined with techniques such as statistical fine-mapping for multiple height-related traits could improve both causal signal identification and understanding of associations with height and height-related phenotypes. Furthermore, fine-mapping has not been performed to date on height-related derived traits, which could help further define the consequences of causal variants, especially in the scenario where there are multiple signals with distinct phenotypic consequences in the same loci (Bartell et al. 2020; Hernández et al. 2021).

The vast majority of genetic research has focused on European ancestry individuals, but GWAS in non-European ancestries are now increasing in size (Nagai et al. 2017; Chen et al. 2011; Loïc Yengo et al. 2022; Conti et al. 2021; Mahajan et al. 2022). Although there is likely to be substantial shared biology across ancestries (Brown et al. 2016), expanding the range of studies in non-European ancestries is important not only to improve predictive power in non-European-ancestry populations but also to leverage signals from multiple populations to better define the location and population-specificity of causal variants (Kanai et al. 2021). In previous work (Kanai et al. 2021), height-associated loci have been fine-mapped in both European and non-European ancestries. However, the presence of multiple signals at each locus, combined with differing LD patterns across ancestries, makes it challenging to define if and where there are truly signals with ancestry-differing effects.

Here, we perform GWAS and meta-analysis of several height-related traits – SHR, leg length, and sitting height – in large biobanks from the UK (UK Biobank, UKB) and China (China Kadoorie Biobank, CKB). Even after correcting for winner’s curse, discovery effect sizes of variants identified in one biobank remained systematically higher than the corresponding estimates from the other Biobank, likely due to differences in their LD with causal variants across ancestries. We perform fine-mapping in UKB, identifying 95% Credible Sets (CSs) implicated in height-derived traits, and identify CSs unique to UKB and not replicated in CKB. Together, these approaches provide additional insights into the genetics and biology of skeletal growth and model a set of approaches for evaluating GWAS from groups of related phenotypes and multiple ancestries.

## Results

### GWAS of skeletal growth phenotypes

We have previously performed a GWAS meta-analysis of sitting height ratio (Chan et al. 2015)SHR, Figure 1A, (Chan et al. 2015) in 21,590 subjects of European ancestry. To uncover the full genetic spectrum of SHR, we extended this study to include 451,921 individuals of European ancestry from the UK Biobank. A deeper understanding of the genetic architecture of SHR can help elucidate the genetics and biology underlying height, skeletal growth, and skeletal growth disorders.

**Figure 1.**
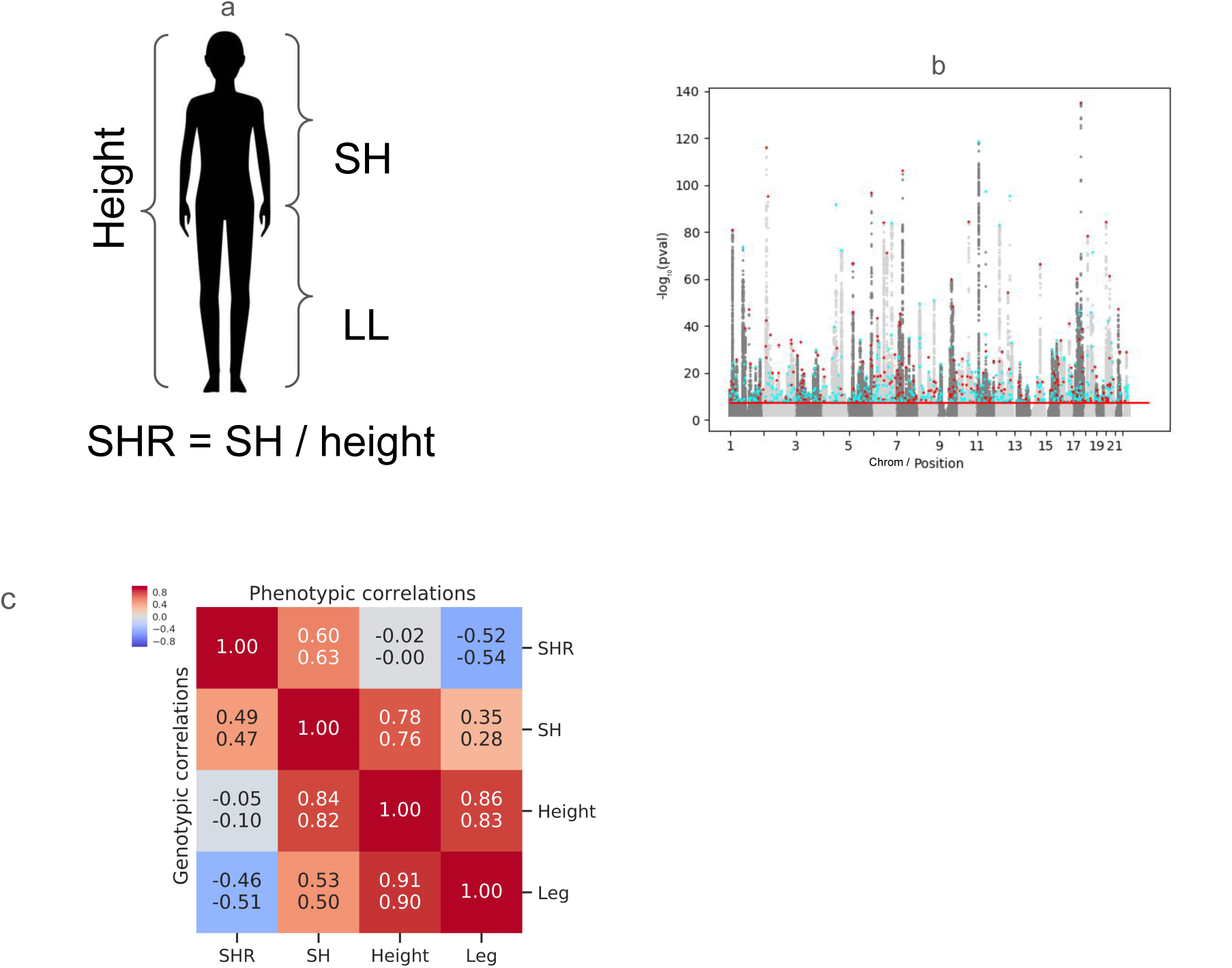
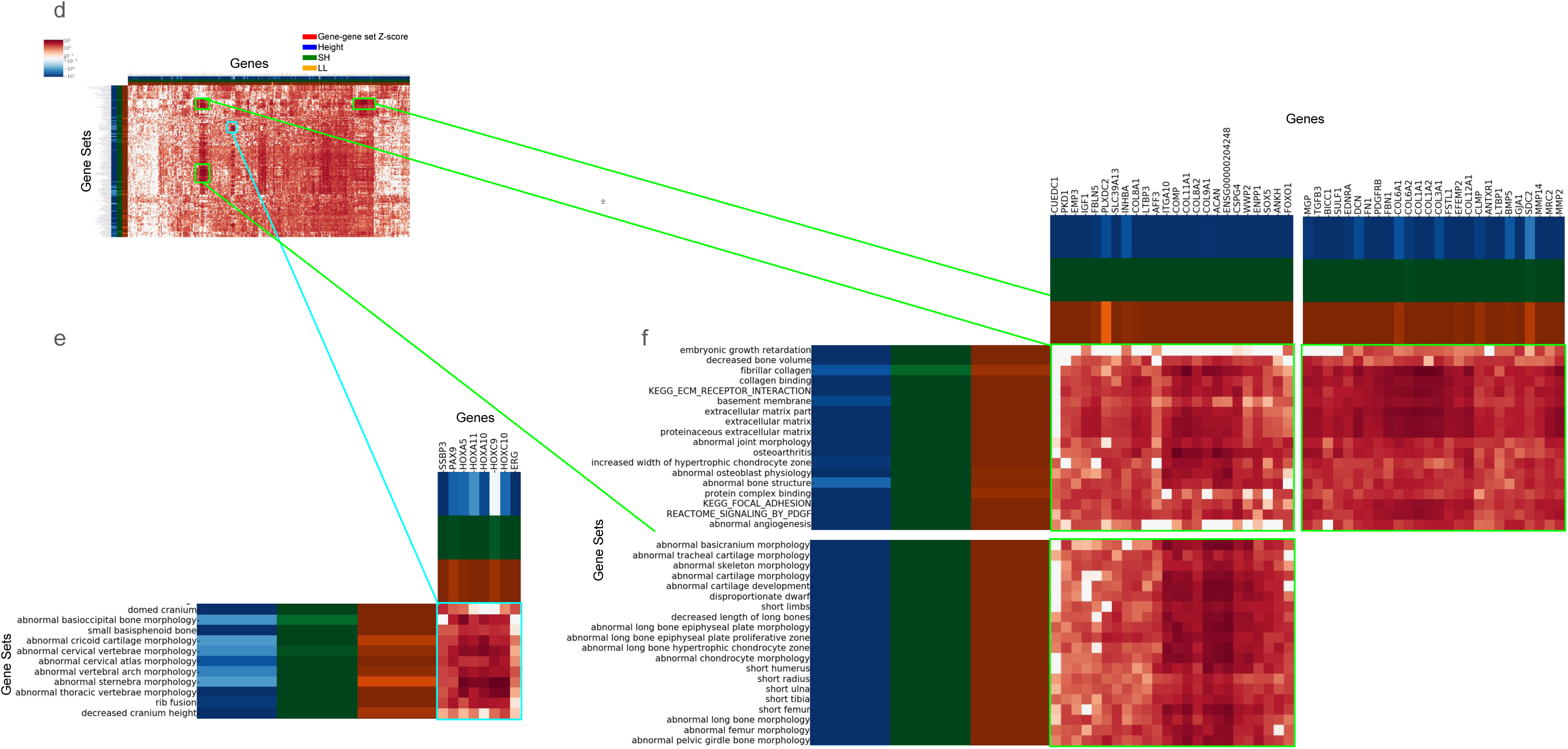
A. Schematic illustrating the phenotypes studied in this manuscript, including height, sitting height (SH), leg length (LL), and sitting-height ratio (SHR), which is SH/height; for most analyses, SHR is adjusted for height and BMI. B. Manhattan plot of the meta-analysis of GWAS results for SHR from UK Biobank (UKB), China Kadoorie Biobank (CKB) and our prior study (Chan 2015). -log10 of p values for association of SNPs with SHR are plotted, with SNPs ordered by chromosome and position. Genome-wide significance (p<5x10-8) is indicated by the dashed line. SNPs reaching genome-wide significance that are also within 35kb of a Height lead SNP (as defined in Yengo 2022) are indicated by red dots and genome-wide significant SNPs falling outside those windows are indicated by blue dots. C. Phenotypic (above the diagonal) and Genotypic (below the diagonal) correlations for height-related traits are show; positive correlations are red and negative correlations are blue. D. Each row is a gene set identified by enrichment analysis (GSEA) of SHR using MAGMA (top 2.5% of prioritized gene sets), and each column is a gene prioritized by PoPs (top 2.5% of prioritized gene sets). The squares in the heatmap indicate the likelihood of membership of the gene in the corresponding gene set (gene-pathway Z-score supplied by the DEPICT reconstituted gene sets Pers 2015). Row and column annotations in the left and top margins indicate whether the gene set or gene, respectively, was also prioritized prioritization by similar analysis of each of the other height-related phenotypes (colored indicates prioritized). E, F. Zoomed regions show clusters of related prioritized genes and gene sets, more prioritized by SHR and similarly prioritized across phenotypes, respectively.

To identify novel loci associated with SHR, we performed genome-wide association in individuals of European ancestry from the UK Biobank (methods for further details). We adjusted SHR for age, BMI, standing height, the genotyping array used by the UK Biobank (UK Biobank Axiom Array or UK BiLEVE), and principal components (PCA) of population structure.

To evaluate phenotypes related to SHR, we performed GWAS of skeletal growth phenotypes sitting height (SH), leg length (LL), and height, as well as SHR unadjusted for height or BMI (SHR-unadjusted) in the UK Biobank. Here, we identified 705 SH, 724 LL, 765 height, 618 SHR, and 643 SHR-unadjusted non-overlapping loci respectively by pruning signals in 500 kilobase pair (Kbp) windows and merging overlapping windows (Figure 1B, Supp Table 2). The observed genomic inflation factor was consistent with previous studies of polygenic traits like height at this sample size, and the LDSC ratio reflects acceptable levels of inflation (P < 5e-8, λ_GC_ = 1.92, Ratio = 0.125 for SHR, Supp Figure 1). Additionally, no evidence of collider bias was observed between height and SHR when comparing SNP effect estimates (R^2^ = 0.01, Supp Figure 2A).

In addition, GWAS of skeletal growth phenotypes were performed in the China Kadoorie Biobank (CKB), using 72,471 unrelated individuals (methods). GWAS was first performed in each of 10 subpopulations and then meta-analyzed. We identified 128, 74, 96, and 57 loci associated with height, SH, LL, and SHR respectively, again using 500Kbp window p-value pruning (P < 5e-8, λ_GC_ = 1.177 for SHR).

### Meta-analysis of UKB and CKB

To identify shared effects between our data sets, a fixed effects meta-analysis of CKB and UKB using METAL (Willer, Li, and Abecasis 2010) was performed (max N = 524392), identifying 732, 672, 686, and 594 loci associated with height, SH, LL, and SHR respectively when limiting to SNPs included in both studies (in turn limiting locus identification).

In addition, prior data (Chan et al. 2015) was included in a separate meta-analysis of SHR. This EUR-specific meta-analysis of 473,511 individuals from 7 studies identified 553 loci at genome-wide significance. Of the five loci that reached genome-wide significance in the previous GWAS of European-ancestry individuals, four were genome-wide significant in UKB and replicated in the meta-analysis (rs6931421, rs1722141, rs882367, and rs228836); one was not significant in UKB (rs140449984, P = 4.1e-3) and therefore showed only suggestive statistical evidence in the meta-analysis (P = 2.468e-5). The association at SNP rs140449984 was primarily driven by one of the six cohorts in the previous GWAS (ARIC).

Finally, an all-sample multi-ancestry meta-analysis of SHR including prior data, UKB, and CKB (total N = 545,984) was performed, identifying 565 non-overlapping loci reaching genome-wide significance when limiting to SNPs present in all studies (Figure 1B; P < 5e-8, λ_GC_ = 1.31, Supp Figure 1).

### Genetic and phenotypic comparison of skeletal growth phenotypes

We sought to estimate the heritability of our SHR, and evaluate the shared genetic component between our phenotypes. To this end, we performed linkage-disequilibrium score regression (LDSC, (B. K. Bulik-Sullivan et al. 2015)) to estimate the heritability of SHR. The variance explained by autosomal SNPs from the 473,511 individuals of European ancestry is 0.22 (Supp Figure 3). We also tested for shared heritability between SHR, height, and other skeletal growth phenotypes. The estimated genetic correlation (B. Bulik-Sullivan et al. 2015) between height and SHR-unadjusted is -0.39 (SE: 0.027) and highly significant (P = 2.31e -46), whereas the estimated correlation between height and SHR (adjusted for height and BMI) is -0.032 (SE = 0.0347) and not significant (P = 0.35), and only slightly negative when estimated in CKB (rg = -0.0999, SE = 0.0404), suggesting that this adjusted phenotype provides a genetically independent assessment of skeletal growth. Other relationships between skeletal growth traits are shown in Figure 1C, and heritability estimates in both ancestries and the meta-analysis can be found in Supplemental Figure 2. Our further analyses below focus on the analysis of SHR adjusted for height and BMI (SHR) and not SHR-unadjusted.

To further address the relationship between derived phenotypes and height, we evaluated the proximity of lead SNPs for different traits. SNPs associated with each phenotype (SHR, SH, and LL) were significantly more likely than null SNPs (matched using SNPSnap (Pers, Timshel, and Hirschhorn 2015), see methods) to be near height signals (p < 0.001, Figure 2A), as may be expected for SNPs associated with height-related phenotypes, and SNP associations with SHR were frequently more significant than with height at a particular height-associated locus (Supp Figure 9). However, even when limiting to associations near height signals, 24% of associations have low LD with the nearby height signals (r^2^ < 0.3 with height lead SNP, within 10Kbp, Supp Figure 10), implying that height-associated regions harbor independent associations with SHR and other height-related phenotypes. Additionally, SNPs associated with SHR-unadjusted fell near height associations more often than SHR (with adjustments for height and BMI); this result argues that the proximity of SHR signals and height signals is not due to collider bias (Supp Figure 2B-C).

**Figure 2.**
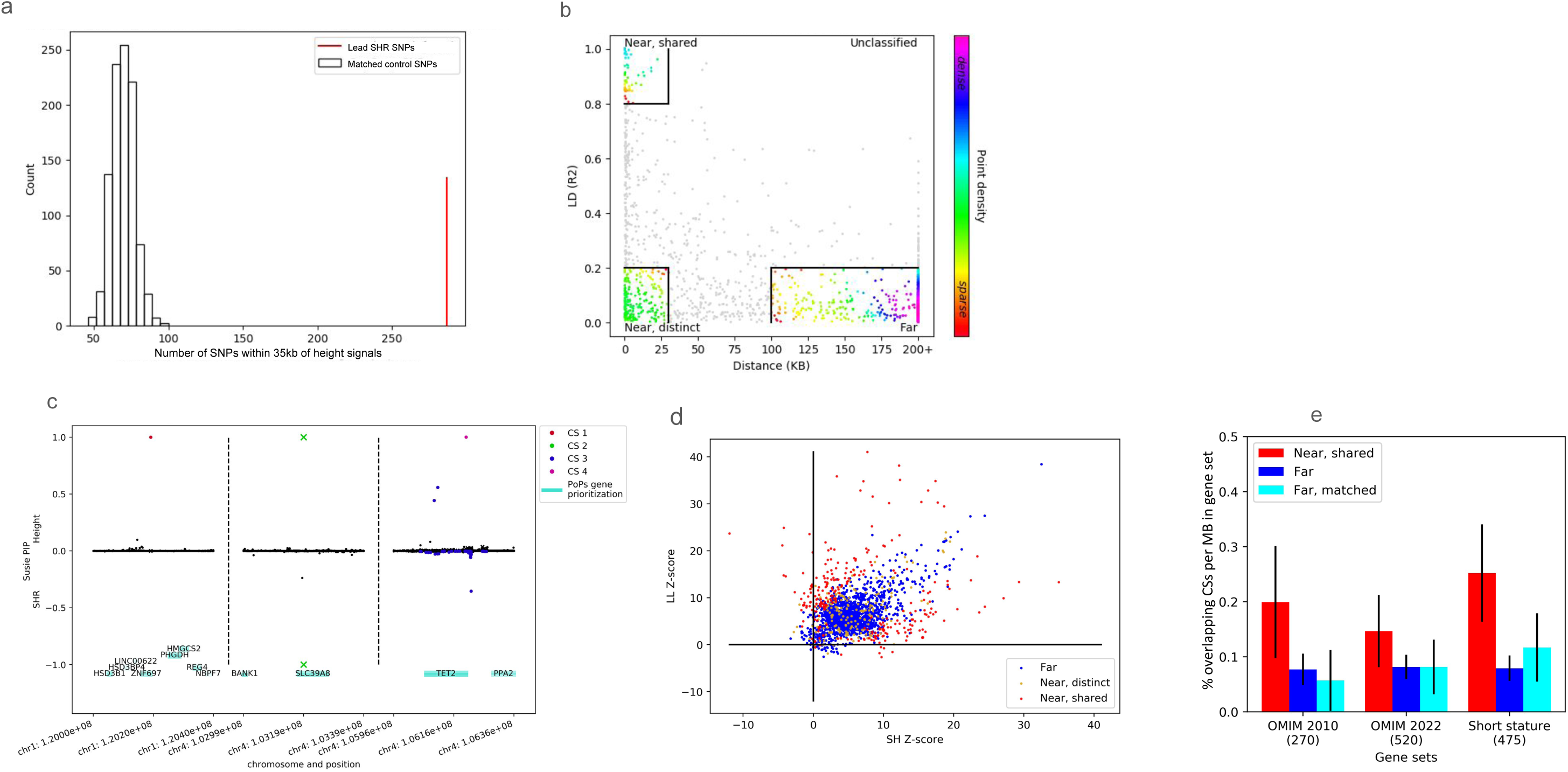
A. Number of lead SNPs (red bar) within 35 kb of height signals of association (COJO lead variants from Yengo et al.), compared with distribution of 1000 control sets of random SNPs matched using SNPsnap (histogram). B. Relationship between height and SHR credible sets (CSs) are plotted with distance (bp) on the X axis and LD (r^2^) on the Y axis. Each point corresponds to one height CS. For CSs containing more than one SNP, minimum distance and maximum LD between SNPs in the height CS and nearby SHR CSs are plotted. Colors mark point density. CSs are classified by their proximity as “near, shared” (282 in top left box, LD between CSs > 0.8 and distance < 30kb), “near, distinct” (189 in bottom left box, LD < 0.2 and distance < 30kb), or “far” (1566 in bottom right box, LD < 0.2 and distance > 100kb); the remainder are “unclassified” (525). C. Examples of height CSs that are “far” (left), “near, shared” (middle), and “near, distinct” (right) from SHR CSs. Colors mark SNP inclusion in a credible set, and X marks a SNP in a credible set for both phenotypes. Regional genes are annotated. D. For height signals proximal to SHR signals, associations with SH and LL are plotted, with SH Z-score on the X-axis and LL Z-score on the Y-axis. “Near, shared” CSs are plotted in red; “near, distinct” are in black; “far” are in blue. 260 / 282 “near, shared” signals maintain significance for association with SHR after Bonferroni correction. E. Overlap between the regions spanned by two different categories of height CSs (“near, shared”, red; “far”, blue) and categories of genes underlying skeletal growth disorders are shown. Black bars indicate 95% confidence intervals. OMIM 2010: (manually annotated list of genes implicated in short stature derived from the OMIM database from Allen 2010); OMIM 2010: (manually annotated list of genes implicated in short stature derived from the OMIM database from Yengo 2022); Short stature: (growth disorders and skeletal dysplasias panel used for genetic testing from Blueprint Genetics). Cyan bars represent “far” CSs matched to “near, shared” CSs, see methods.

### Gene Set Enrichment Analyses

To explore genes and biological pathways and genes underlying SHR, gene set enrichment analyses (GSEA) were performed using MAGMA (de Leeuw et al. 2015) to meta-analyze GWAS summary statistics from UKB and CKB. In line with observational and genetic correlation results, height and LL gene-set significance was qualitatively most similar, both identifying height-LL prioritized pathways (“cell cortex”, “cell leading edge”) and pathways shared with all 4 phenotypes (“ossification”, “decreased length of the long bones”) (Supp Figure 4); height and SHR gene set significance was qualitatively most different (Supp Figure 4). SHR gene-set enrichment with MAGMA identifies growth-related pathways (e.g. “embryonic growth retardation”, “short tibia”). We also applied PoPs (Weeks et al. 2020), which prioritizes genes based on similarity to genome-wide enrichment, to prioritize genes related to SHR. PoPs identifies growth-related genes (e.g. *GHR*, *COL1A1*, *ACAN*) (Figure 1D). We also sought to identify biology affecting skeletal proportion but specifically not height; we observed that a cluster of *HOX* genes was more prioritized for SHR and SH than for height or LL (Supp Figure 5) and were prioritized for pathways relating to vertebral morphology and branching morphogenesis. When analyzing genes selectively prioritized by SHR but not height, Gene Ontology (GO) identifies “negative regulation of animal organ morphogenesis” (GO:0110111) and “negative regulation of Wnt signaling pathway involved in dorsal/ventral axis specification” (GO:2000054) among other relevant pathways (prioritized list in Supp table 7).

### Fine-mapping of signals in UKB

Since the above analyses were limited to lead signals per region, they do not conclusively demonstrate separate effects on related phenotypes in complex loci; to address this, we performed statistically-informed fine-mapping. Due to limitations of existing fine-mapping methods that have been shown to result in substantial miscalibration when applied to meta-analyses (Kanai et al. 2022), fine-mapping was performed using SuSiE (Wang et al. 2020), Methods) on UKB GWAS summary statistics. Overlapping loci (where lead SNPs were within 1 megabase pair (Mbp) of each other) were merged, resulting in 765 height-associated, 618 SHR-associated, 705 SH-associated, and 724 LL-associated loci with average sizes of 2.81, 2.19, 2.5, and 2.5 Mbp respectively. To computationally enable fine-mapping even at >1Mbp loci containing many SNPs, analyses were limited to a subset of SNPs per locus (those with P < 0.01 or identified as part of a CS in a 1Mbp analysis, see methods, Supp Figure 5). For height, there were on average 79,951 SNPs per variable-sized locus, of which 18,487 per locus were considered in the fine-mapping model. Across all loci, an average of 73 SNPs were members of 3.8 credible sets (CSs) per locus (Supp Figure 7). A total of 2817 CSs were identified containing 47,740 SNPs, of which 1,537 contained 5 or fewer SNPs. The majority of loci (482) contained 5 or fewer CSs. These and the corresponding statistics for fine-mapping of each of the other traits can be found in Supp Table 1, and full SNP-level summary statistics can be found in Supp Table 6.

### Comparing effect sizes of fine-mapped loci across phenotypes

Credible sets identified for each phenotype were evaluated based on their proximity to or overlap with CSs identified for other phenotypes. As shown in Figure 2B, some height CSs were identified both in close proximity to and (sometimes) overlapping SHR CSs, while other height CSs were relatively distant from SHR CSs in both genomic distance and LD. Using these metrics, we classified groups of CSs: “near, shared” if the minimum distance was less than 30Kbp and the maximum LD r^2^ was greater than 0.8 between members of credible sets from phenotypic pairs (282 CSs); “near, distinct” if the distance was less than 30Kbp but the maximum LD was r^2^ < 0.2 (189 CSs); “far” if the distance was greater than 100Kbp and the maximum LD was r^2^ < 0.2 (1566 CSs); and “unclassified” otherwise (525 CSs). In a specific example of a “near, shared” on chromosome 4 (Figure 2C), a credible set overlapping the *SLC39A8* gene is shared between height, SHR, SH, and LL; this specific CS associates with opposite-direction effects on SH and LL. The allele of the single-SNP CS associated with increasing height associates with increasing LL but decreasing sitting height, leading to a decreased SHR. Several further example loci are provided in Supp Figure 14. As expected, most height CSs “near, shared” with SHR are estimated to affect SH and LL differently or have opposite-direction effects on SH and LL (Figure 2D, Supp Figure 11, methods).

We then sought to characterize the proximity of CSs to height-relevant genes. To do this, we used manually curated gene lists: first, height-related genes from the Online Mendelian Inheritance in Man (OMIM) database (Lui et al. 2012) as described in (Lango Allen et al. 2010) (“OMIM 2010”) and (Yengo et al. 2022) (“OMIM 2022”), and second, short-stature causing genes (“Genetic Testing for Growth Related and Skeletal Disorders” 2021). Height CSs “near, shared” to SHR CSs more often overlapped height relevant genes than height CSs “far” from SHR CSs (Figure 2E); this conclusion remains even when subsetting “far” CSs to match “near, shared” GWAS association p-values with height. These short-stature causing genes were further classified as causing syndromes of short stature proportionally to upper and lower body (e.g. fanconi anemia, (dos Santos, Gavish, and Buchwald 1994)) or disproportionally (e.g. spondyloepimetaphyseal dysplasia, (Cormier-Daire 2008)) (see methods, Supp Table 5). Because SHR is a measure of skeletal proportion, we then asked whether the overlap with short stature was driven by genes underlying syndromes with disproportionate short stature. We compared the proportion of genes that overlapped “near shared” and far signals in these 2 groups of short stature genes. As might be expected, fewer of the “far” CSs overlap disproportionate genes than proportionate genes compared with the “near, shared” CSs; however, this observation was not significant so further work would be needed as confirmation (D^2^ test, p > 0.05; Supp Figure 12).

Our previous work had shown that genes implicated by height GWAS showed expression specificity for the round layer of the growth plate (Renthal et al. 2021). We sought to test if this observation replicated among our fine-mapped CSs. Genes overlapping height CSs “near, shared” to SHR CSs and height CSs “far” from SHR CSs both had higher round zone expression specificity than all assayed genes (t-test P = 2.1e-13, 3.2e-8 respectively), and these “near, shared” CSs additionally had significantly higher round zone specificity than the “far” CSs (P = 2.4e-4, Supp Figure 13), consistent with the prior results (Renthal et al. 2021) and emphasizing the biological relevance of genes overlapping “near, shared” CSs. Identification of these loci provides specific targets for further interrogation of the biology of human growth.

### Heritability and genetic correlation in multiple ancestries

Phenotypes observed in UKB are similar between ancestral groups, but ancestry-specific averages remain statistically significantly different ((Eveleth and Tanner 1991; Bogin and Varela-Silva 2010; Chan et al. 2015); UKB EAS vs EUR SHR P = 4e-97, Supp Figure 17). Therefore, we sought to explore any potential genetic basis for such observed differences. LDSC was used to calculate heritability of each phenotype in CKB, showing lower heritability in each phenotype. In addition, we estimated heritability using GWAS of each phenotype performed in UKB, downsampled to match CKB (Supp Figure 3). For each phenotype, genetic correlations between ancestries were then evaluated using Popcorn ((Brown et al. 2016), see methods). Each phenotype’s cross-ancestry genetic correlation was not significantly different from 1 (Figure 3A).

**Figure 3.**
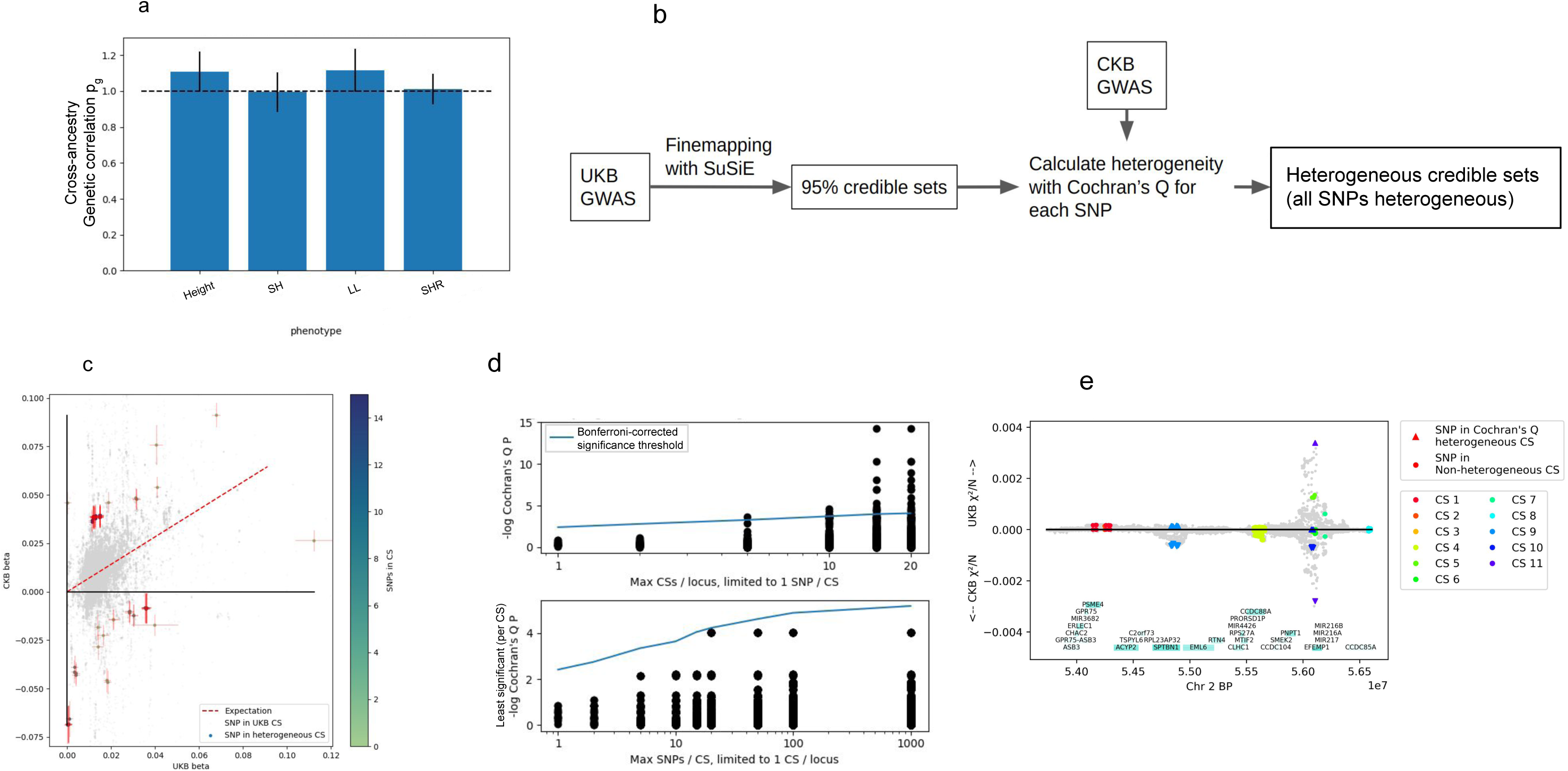
A. Cross-ancestry genetic correlation as evaluated using Popcorn between GWAS derived from UKB and CKB across phenotypes height, SH, LL, and SHR adjusted for height and BMI. Estimates for each phenotype are not significantly different than 1. B. Schematic of heterogeneous CS identification. C. Heterogeneous CSs identified for height. UKB GWAS is finemapped using SuSiE, and effect estimates for SNPs in credible sets were compared with estimates from CKB using Cochran’s Q. CSs that were entirely composed of heterogeneous SNPs were nominated as heterogeneous credible sets. For each CS, the least heterogeneous SNP is plotted. D. For height, limiting to most simple CSs (max CSs per locus = 1 (top), max SNPs per CS = 1 (bottom)), and then relaxing that both limits, heterogeneity significance of CSs as multiple testing threshold increases. Cross-ancestry heterogeneous CSs were only observed in loci with multiple credible sets. E. Locus where finemapped SNP effects differ between CKB and UKB. SNPs in credible sets are circled, and SNPs in the 3 heterogeneous credible sets are marked with triangles.

### Comparing effect sizes of lead signals across ancestries

To evaluate the extent of shared genetic effects on height between ancestral groups, we first investigated lead SNPs from both UKB and CKB GWAS. Effect estimates in a discovery dataset tend to be overestimated due to the effects of Winner’s Curse, or in effect choosing SNPs that performed well in this analysis such that they were called genome-wide significant when in fact their true effect size would not be genome-wide significant in this study. As such, we analytically corrected the discovery effect sizes for Winner’s Curse (WC) (Palmer and Pe’er 2017; Zhong and Prentice 2008), and then evaluated systematic differences in effect size between populations using the difference of means test. First, to test the efficacy of Winner’s Curse correction, lead SNPs were identified for skeletal growth phenotypes from GWAS using down-sampled unrelated subsets of half of UKB or CKB as the discovery population. The second unrelated half of the discovery population was used as an unbiased control set, and GWAS from the second population used to evaluate differences between populations. Winner’s curse corrected GWAS from half of UKB compared with CKB showed a significantly larger effect size difference than when comparing with an equal sized independent subset of UKB (t-test, P = 3e-18 for height). Notably, although the direction of effect is the same, the reverse analysis (comparing corrected CKB GWAS against half of UKB, versus with an independent subset of UKB compared with the half of UKB) is not significantly different; this may be due to lower power in association identification and a consequently larger correction for WC. WC is controlled through correction, albeit potentially over-corrected (Supp Figure 15), and the corrected data set acts similarly to using a replication data set (Supp Figure 15). Full GWASs were compared across each phenotype (Supp Figure 16), effect sizes are larger in the discovery population despite WC correction. Each of the 4 skeletal growth traits showed a similar amount of evidence for population differences (Supp Figure 16J).

### Comparing effect sizes of fine-mapped loci across ancestries

To identify examples of height associations where observed effect sizes in UKB differed from those in CKB, fine-mapped CSs were used. UKB credible sets were identified as “ancestrally heterogeneous” (A-Het) if all SNPs in the credible set showed significant heterogeneity of effect size between UKB and CKB (described in schematic 3B). CSs displaying A-Het were concluded to have differential effects in UKB and CKB (Figure 3C, Supp Figure 18). Of 8554 credible sets identified across 4 phenotypes in UKB, 53 CSs in 44 loci contained only A-Het SNPs (P < 0.05 / # total CSs per phenotype) and 9 loci contained more than one A-Het CS. Additionally, 9 A-Het CSs were shared between phenotypes, one of which was shared between 3 phenotypes: height, SH, and LL. Compared to CSs that included SNPs that were not A-Het, A-Het CSs had fewer SNPs per CS and were located in loci with more CSs; SNPs in A-Het CSs were more common than the full set of SNPs in all CSs. Examples of cross-ancestry heterogeneity are provided in Supp Figure 19.

When restricting these analyses to the 14 SHR-associated loci containing a single CS comprising a single SNP, only 1 significantly A-Het locus was identified despite partially alleviating the multiple-testing burden. This single SNP CS (3:188365278:A:G, rs4686991) is associated with SHR in UKB and CKB in opposite directions (CKB marginal beta = 0.0401 (SE = 0.0204, P = 0.0496) vs UKB marginal beta = -0.024792 (SE = 0.003156, P = 4.8e-16)), and lies within an intron for the *LPP* gene which encodes LPP, a LIM domain containing protein that may have roles in cell-cell adhesion or cell motility (Kuriyama et al. 2016); the CS also falls ∼41Kbp upstream of MIR28, a miRNA involved in cell proliferation (Schneider et al. 2014). Relaxing filtering for any of the phenotypes to allow more than 1 CS in a locus led to identification of further A-Het CSs, but not when allowing for more than 1 SNP in a CS (Figure 3D). In total, we identified a finite number of CSs showing differences in SNP effects between ancestries, only one of which is a single-SNP CSs and few of which are in simple loci with few CSs.

## Discussion

The familial basis of human skeletal growth has been the subject of study since before the advent of human genetics (Galton 1886), but new datasets and methods continue to provide new insights. The largest GWAS to date, of human height, has identified over 12,000 approximately independent height-associated signals (Loïc Yengo et al. 2022). This study mapped essentially all of the regions where common variation contributes to heritability, and additionally found that current approaches to prioritize effector genes and gene sets saturate at these larger sample sizes. Despite the progress in recent work, however, it remains an open question as to how best to translate these findings to specific insights about the biology driving human growth. Past work has used computational approaches or functional data in relevant models to provide more insight in the context of height (Bartell et al. 2020; Muthuirulan and Capellini 2019; Baronas et al. 2023; Wood et al. 2014) as well as for many other phenotypes (Welter et al. 2014; Lappalainen and MacArthur 2021; Amariuta et al. 2019), but further development of methodology in addition to application to larger and more ancestrally diverse datasets are necessitated for future advancement.

Here, we use additional genetic association datasets - specifically, GWAS of height-related phenotypes - as a tool to understand the biology driving height. Past work involving GWAS of multiple related phenotypes has explored improving power for genetic discovery (Turley et al. 2018), identifying loci whose effects overlap between phenotypes (Shi et al. 2017; Giambartolomei et al. 2014), or classifying loci based on their effect on related phenotypes (Udler et al. 2018). These approaches, however, may not be best-equipped to handle complex loci containing many signals, as is often the case for loci associated with polygenic phenotypes with large sample sizes, such as height. In this work, we focused on disentangling putative causal signals within loci, with the goal of distinguishing nearby but independent effects, allowing us to identify loci where nearby signals have distinct consequences on height-related phenotypes.

Our work increased the number of loci found to be associated with SHR from 6 (in our earlier study) to 565. From these data, we prioritized biologically relevant genes and gene sets, and fine-mapped signals specific to or shared between height-related phenotypes. By comparing results from two major ancestries, we could describe individual SNP-level differences in effect sizes and identify loci where effect sizes are consistently different across credible sets of potentially causal variants; these loci comprise a small minority of associated loci.

Leveraging associations with each of four height-related phenotypes, we could evaluate the shared and distinct components of each. Height-associated loci are observed both near and far from associations with each of the other three phenotypes, with LL-associated loci showing the greatest proximity. Additionally, height-associated signals identified in Yengo et al. 2022 are often located near loci with more significant associations with SHR, SH, or LL than height, despite being ascertained on height, which has greater power for detection due to higher heritability in UKB. This emphasizes the potential value of these height-related traits in understanding underlying skeletal growth biology. Height is highly genetically and phenotypically correlated with SH and with LL, but not with SHR (when corrected for height), which is in turn moderately correlated with SH and LL, as expected. As expected given the variability in SHR, SH and LL are only moderately correlated. Gene set enrichment analysis identifies gene sets primarily related to growth and structural development (e.g. limb development, collagen binding), as well as general biology (e.g. transcription factor binding) for each of the four traits. In addition, well-known genes related to growth were prioritized, and gene and gene set prioritization for each trait can be found in supplemental tables (Supp Tables 3, 4). SHR GSEA prioritized mostly genes and pathways shared with the other phenotypes but in particular *HOX* genes (which are known to be implicated in development of body structures, (Hajirnis and Mishra 2021)) were prioritized more strongly for SHR than height.

We report here fine-mapping results for loci associated with each of the 4 phenotypes, and identify many SNPs marked as being a part of a credible set for more than one trait. In addition to expected shared signals between related phenotypes, we also identify many credible sets specific to a single phenotype. Notably, credible sets near and shared between height and SHR, compared to height-only credible sets, were enriched for multiple metrics of relevance to the biology of growth.

Our previous study (Chan et al. 2015) evaluating the genetics of SHR demonstrated its high heritability and its potential use as a tool to understand height-associated variants. Here, we meta-analyzed these results with GWAS using the UK and China Kadoorie Biobanks, increasing total sample size from 25,135 to 545,984. The associated increase in statistical power allows use of multiple downstream methods to investigate each phenotype, including evaluation of predictive polygenic risk scores, gene set enrichment, and fine-mapping of individual loci. Additionally, we perform GWASs of sitting height and leg length, which are each closely related but distinct from height.

In addition to increasing power, including multiple ancestries also allowed us to ask questions about differences in putatively causal loci and their effect sizes across ancestral groups. Using GWAS across two ancestries, we looked to identify similarity in effect sizes genome-wide. After validating and applying correction for winner’s curse, we observed effect sizes were on average larger in the discovery dataset. Because the underlying genes and pathways are known to be broadly conserved between ancestries, this genome-wide difference in effect size is most likely due to the fact that GWAS lead SNPs often only tag the causal signal in a region but are not themselves causal (Visscher et al. 2012). When comparing effects between cohorts with different continental ancestries, there may be local differences in LD such that the lead SNP in one ancestry does not strongly tag the true causal variant in the other ancestry, leading to an observed decrease in both significance and effect size. In order to better understand examples of differences in effect, an approach that takes LD into account is needed when comparing effect sizes across ancestries.

To begin to explore this, we performed fine-mapping (in the better-powered European GWAS), nominating credible sets (CSs) where we are reasonably confident (alpha>0.95) that the CS of SNPs contains a truly causal SNP. If we observe heterogeneity in effect size between ancestries at all the potentially causal SNPs in the CS, we can be confident that differing LD patterns are no longer a potential explanation for the change in effect size. We identified a small proportion of CSs (0.6% for height) with heterogeneous effect sizes across ancestries for all SNPs in the CS. Although it is possible that the same causal SNPs are present in each ancestry but have different effect sizes due to differing genetic background (potentially due to SNP-SNP interactions, which remain challenging to detect (Westerman et al. 2022)), or that there is an ancestry-specific causal variant, it is also possible that these observed differences in effect are due to interactions with population-specific environmental factors, or differences in sample ascertainment. Of note, the CSs showing complete heterogeneity of effect size were mostly seen in loci with many different signals of association, raising the possibility that some of this heterogeneity could be explained by insufficiently precise fine-mapping at these complex loci, leading to the actual causal SNP not actually being in the CS.

Our work based on fine-mapping is subject to additional limitations, some of which we discuss here. Fine-mapping can be sensitive to which SNPs are included in the model (Kanai et al. 2021; Ulirsch 2022), and our computational approach to performing fine-mapping of regions larger than 1Mbp limited the number of SNPs included in the model to a subset of SNPs more likely to be causal. However, our comparisons across both phenotypes and populations remain similar when only considering loci of size 1Mbp, implying that our specific implementation for larger loci did not substantially bias the findings. It is also possible that some causal variants are not in our dataset (due to low frequency, type of variant, and/or filtering based on poor imputation/missingness), resulting in a CS incorrectly being designated as heterogeneous even though the missing causal variant has consistent effect sizes across ancestries. Additionally, this study only describes fine-mapping analyses performed on European ancestry GWAS. Future work must be done to extend this approach to populations with additional ancestries, especially as sample sizes continue to grow, in order to ensure that findings from this and other genetic studies are equitable for all people. In particular, because of differences in available sample sizes, our approach is only powered to identify heterogeneity of effect sizes at loci discovered in UKB, as opposed to loci identified specifically in CKB. For instance, we chose not to analyze the approximately 2364 individuals with East Asian ancestry in the UK biobank to preserve homogeneity of individual GWAS analyses (Constantinescu 2022), and further method development would be needed to best integrate these samples to maximize equity of discovery. Finally, our approach, which requires heterogeneity at all SNPs in a CS, is likely biased to identify heterogeneity of effect size in small rather than larger CSs.

In summary, we have explored the genetics of height-related skeletal growth phenotypes – SHR, SH, and LL, with substantially increased power to identify genetic associations with these phenotypes. We identify shared biology across phenotypes through GSEA, prioritizing relevant genes and pathways. We additionally identify shared and distinct genetic effects between phenotypes through the use of statistical fine-mapping and use this to categorize height-associated variants into different phenotypic categories that have different biological properties. We also propose a preliminary fine-mapping-based approach to identify loci where effects at causal SNPs are likely to differ between ancestries. As sample sizes continue to increase and fine-mapping methods continue to improve, these and similar approaches will enable a more thorough dissection of causal genes and variants, their phenotypic consequences, and any differences in effect sizes across ancestries. Taken together, this work illustrates advantages of integrating genetic association data across related polygenic phenotypes and multiple ancestries to better understand underlying genetic and biological mechanisms.

## Materials and Methods

### GWAS in UKB

GWAS was performed on phenotypes height, sitting height (SH), leg length (LL, height - SH), and sitting height ratio (SHR, SH / height) using 206812 males and 245109 females of European ancestry from the UK Biobank over 77753733 autosomal SNPs passing quality control filters. Phenotypes were adjusted for Age and Sex (for joint analyses of males and females). Sitting height ratio GWAS was performed in multiple ways, either with the same covariates as other phenotypes or additionally correcting for BMI or height and BMI. Genotyping array number and genetic principal component were additionally included as covariates in GWAS. GWAS was performed using Bolt-LMM version 2.3.4. Additionally, GWAS of independent halves of UKB (N = 188575, 188597), UKB size-matched with CKB (N = 72340, 72321) subset from the halves, and thirds of UKB (N = 125740, 125735, 125748) was performed and was used in supplemental PRS analyses.

### SHR EUR Meta-Analysis

UK Biobank GWAS results were meta-analyzed with previously published SHR genome-wide association data. The previous GWAS of SHR used data from six studies, for a combined sample size of 21590 individuals of European ancestry and 3545 African Americans. Imputation for all six cohorts was performed using a 1000 Genomes reference panel, and only SNPs with an information score > 0.6 were included in the meta-analysis. From the UK Biobank study, SNPs with an information score > 0.3 were used. METAL version 2011-03-25 (Willer, Li, and Abecasis 2010) was used.

### GWAS in CKB

China Kadoorie Biobank (CKB, https://www.ckbiobank.org) is a population-based prospective cohort. The blood samples of the adult subjects were collected from 10 geographically defined regions of China in 2004-8. Hard-genotyping was conducted on custom Affymetrix Axiom® arrays, with 511885 variants passing QC. Genotypes were phased using SHAPEIT3 v4.12 and imputed into the 1000 Genomes Phase 3 reference with IMPUTE4 v4.r265 as described previously (Walters et al. 2022).

The four traits are standing height, sitting height, leg length (standing height – sitting height) and SHR (sitting height / standing height). Across the 20 region by gender strata, SHR was linearly regressed against age, squared age, year of birth (as a categorical variable), height and BMI. The three other traits were regressed against age and squared age. The residues were rank-based inverse normal transformed and used for GWAS.

The CKB genotype data were split into 5 different sets. The set A is the full set of 95674 subjects. The set B is the relative-free subset of the set A. The individuals were selected via the command ‘plink2 –king-cutoff 0.05’. The set has 72471 subjects. The set C is the relative-free subset of the set A- B. Again, the samples were selected via command ‘plink2 –king-cutoff 0.05’. It has 15620 subjects. The set D is a randomly selected subset of the set B. Its size is 36235. The set E is the set B – D, of 36236 subjects.

We used BOLT-LMM v2.3.2 for the GWAS of the transformed residues. The covariates were the Genotype array version and the sample disease ascertainments. The fix-effects meta-analysis across the 20 region by gender strata was performed using METAL.

### Trans-ethnic fixed-effects meta-analysis

Fixed-effects meta-analysis was performed between GWASs performed in UKB and CKB using METAL. SNPs included in only 1 of the 2 populations were included in reported summary statistics. 473511 EUR subjects and 72473 CKB subjects were combined for a total sample size of 545984.

### GSEA in each population

MAGMA was performed using GWAS summary statistics calculated in UKB and separately in CKB based on reconstituted gene sets (DEPICT (Pers et al. 2015)). A Bonferroni-corrected significance threshold (corrected for number of gene sets) was used to identify significant gene sets at p < 0.05/14462. 1k Genomes was used as the reference panel for gene significance, using EAS and EUR ancestries for CKB and UKB respectively. MAGMA version v1.09 was used.

### GWAS subsets

GWASs of multiple sizes were performed as shown in this table.

| Name | Description | N (height) |
| --- | --- | --- |
| UKB | EUR ancestry in UKB | 453925 |
| UKB unrelateds | EUR UKB without related individuals | 377117 |
| UKB half 1 | A random sample of half of UKB unrelateds | 188575 |
| UKB half 2 | The complement of UKB half 1 from UKB unrelateds | 188597 |
| UKB ckbN half 1, 2 | A random sample of the same size of CKB, taken from UKB half 1 or half 2 | 72340, 72321 |
| UKB third 1, 2, 3 | A random Sample of independent thirds of UKB unrelateds | 125740, 125735, 125748 |
| UKB two thirds | Complements of UKB thirds 1,2,3 | 251422 |
| CKB unrelateds | Unrelated subset of CKB | 72473 |
| CKB half 1, 2 | A random sample of half of CKB unrelateds, complementary | 36237, 36236 |
| CKB held out set | CKB - CKB unrelateds | 15620 |

### Lead SNP proximity

The proximity of lead SNPs identified for each phenotype was compared. A null distribution was generated by matching each lead SNP with up to 1000 null SNPs using the SNPSnap software (Pers, Timshel, and Hirschhorn 2015), matching on default parameters minor allele frequency (+/- 5%), gene density (+/- 50%), distance to nearest gene (+/- 50%), and LD buddies (+/- 50% using r^2^ > 0.5).

### Height-relevant gene sets

Height relevant gene sets from OMIM and from Blueprint Genetics were used to validate fine-mapping results. Height relevant genes from OMIM were manually curated (Loïc Yengo et al. 2022; Lui et al. 2012). The short stature genetic testing panel from Blueprint genetics was used (“Genetic Testing for Growth Related and Skeletal Disorders” 2021), and additionally, genes were classified by their clinical indications as acting in a body region proportionate (changes in height associated with gene disfunction tend to not specifically target upper or lower body) or body region disproportionate (changes in height associated with gene disfunction tend to act majoritarily on one body region). Syndrome implications were classified referring to omim.org.

### Round zone specificity

Mouse gene expression specificity to different parts of the growth plate was calculated as previously described (Renthal et al. 2021). Human orthologues for mouse genes were identified using the biomart package, Ensembl build 105. CS scores for expression specificity were identified for CSs overlapping a gene; averages were used when overlapping multiple genes. Human genes mapping to multiple mouse used averages.

### Winner’s Curse correction

SNPs passing genome-wide significance were identified in summary statistics from GWAS performed in half of UKB or half of CKB; the effect size of these SNPs were shown to be inflated, mostly near the significance threshold when compared to the unrelated second halves (Supp Figure 15). As such, effect sizes were corrected for Winner’s Curse (Palmer and Pe’er 2017; Zhong and Prentice 2008). Optimal correction was achieved when using linear regression P-values from Bolt-lmm. After showing the efficacy of Winner’s Curse correction, the method was applied to the full UKB and CKB GWASs.

### Fine-mapping

Fine-mapping was performed using SuSiE version 0.8.0 on 1MB windows surrounding each lead SNP for unrelated GWAS of phenotypes height, SHR, SH, LL, and SHR-unadjusted measured in UKB. The following parameters were set: maximum number of non-zero effects L = 20, maximum iterations max_iter = 1000, and SuSiE was used to estimate scaled prior and residual variances. LDstore version 2.0 was used to calculate LD in the 1MB windows from European unrelated individuals UKB.

For overlapping 1MB windows, additional analysis was performed. Overlapping windows were merged, and fine-mapping was performed on the entire combined region jointly. In order to save on computational time and space, many SNPs were removed; SNPs were only included in the model if either their p-value for the GWAS of interest was less than 0.01 or if the SNP was identified as being a part of a credible set in the 1MB fine-mapping analysis. As a result, 23.1% of SNPs were included in fine-mapping analyses of merged regions greater than 1MB. A small number of fine-mapping analyses using all SNPs of a merged region were performed, and these were compared to fine-mapping performed with the limited SNP set with high concordance of results (see supplemental text, Supp Figure 6).

### Effect size comparison

Effect sizes at lead SNPs in CKB and UKB were compared. Winner’s Curse corrected trait-increasing effect sizes for UKB and CKB lead SNPs were compared with effect sizes for the SNPs calculated in the second study and their relationship was quantified using Least Squares Regression and Deming Regression. Cochran’s Q and I^2^ were calculated for lead SNPs identified in either UKB or CKB by comparing Winner’s Curse corrected effect sizes with the SNP’s effect size in the second study. I^2^ vs Cochran’s Q P-value are shown in Supp Figure 16H-J.

Fine-mapped CSs were evaluated for ancestral heterogeneity. If all SNPs in a CS were heterogeneous at Bonferroni-corrected significance (Cochran’s Q, p < 0.05 / Credible sets) when comparing betas estimated in UKB and CKB, then the CS was classified as heterogeneous.

### Trans-ancestry genetic correlation

Popcorn (Brown et al. 2016) version 0.9.9 was run on each phenotype to identify trans-ancestry genetic correlations. The “fit” command using pre-computed 1000 genome ld references for each of EUR and EAS ancestral groups was run.

### PRS in UKB and CKB

PRS was performed using the Pruning and Thresholding framework implemented in the PRSice version 2.2.12. PRS was performed for phenotypes height, SHR, SH, LL, and SHR-unadjusted, and performed independently for CKB and for UKB. GWAS from half of individuals from the study was used for effect sizes; individuals from an independent unrelated quarter were used to find the best fit p-threshold and prediction accuracy was evaluated in the final unrelated quarter. SNPs with INFO < 0.3, MAF < 0.005, or ambiguous alleles were filtered.

### Proxy-informed trans-ethnic PRS

Trans-ethnic PRS was created to improve prediction using the Pruning and Thresholding (P+T) framework as implemented in PRSice (Choi and O’Reilly 2019). The following algorithm (shown in Supp Figure 20) was used to generate the list of SNPs used for prediction: GWAS was performed in Study 1 half 1; naive PRS used direct GWAS summary statistics. SNPs were filtered based on MAF and INFO in both studies (half 1 from each study); this list of SNPs was used for initial, filtered, PRS. The SNP list was then clumped by significance, and LD proxies with r^2^ > threshold1 to a lead SNP were identified using Study 1 ancestry. LD proxies with r^2^ > threshold2 to this new list were next identified using Study 2 ancestry. This final SNP list was clumped and PRS generated using p-values from Study 2 half 1. LD information from the UK10K study was used for UKB proxy identification steps, and LD information from 1kg EAS was used for CKB proxy identification steps. RSID identification was performed using hg19 dbSNP 151 (Sherry, Ward, and Sirotkin 1999), and all ambiguous SNPs (A/T or G/C SNPs) were removed from the analysis prior to clumping. For analyses where threshold 2 = 1, the second proxy identification step was skipped; for analyses where threshold 2 = “window”, boundaries were defined using SNPs from the first proxy identification step and all SNPs within these boundaries were included. Independently, for analyses where threshold 1 = “1MB” or other distance, lead SNPs were identified and nearby SNPs within the described window (ie 1MB) were included for Study 2 pruning.

## Supplementary text

### Fine-mapping loci larger than 1MB

As described above, fine-mapping was performed for loci larger than 1MB in size. To make this analysis feasible genome-wide across multiple phenotypes, fewer SNPs were included in the SuSiE model. As implemented, a large portion of time and memory was spent handling the LD matrix, which grows with the square of the number of SNPs, and decreasing the size of this matrix without meaningfully impacting results was of great value to this work. We first performed fine-mapping on 1 MB loci, but noticed where they overlapped, they did not agree. Furthermore, fine-mapping combined loci (using all SNPs, ranging between 1.5 and 2 MB) often gave conflicting results to both of the 1MB analyses. Fine-mapping was performed on a small number of combined loci, but became computationally infeasible when scaling to the full genome. Multiple approaches were taken to remove SNPs deemed unlikely to appear in final credible set predictions; the approach with the best concordance while also substantially improving runtime was identified to be including only SNPs that either 1. passed a significance threshold of p < 0.01 or 2. were identified as appearing in a credible set in 1MB window fine-mapping (Supp Figure 5). This second case was added to include a small number of potentially important SNPs at low cost to runtime. It is likely that some true causal SNPs were removed from the fine-mapping model, and this remains a limitation to our approach.

### Polygenic risk score prediction

Unrelated samples of UKB and CKB were split and subset to create independent populations; GWAS was performed. Using GWAS performed in half 1 of the discovery population and genotype and phenotype data from quarter 3 of the discovery population, Polygenic Risk Scores (PRS) were generated using PRSice, predicting in quarter 3 of the discovery population. In this way, the optimal p-value threshold was identified, and PRS prediction accuracy was then evaluated by again generating PRS (using the same p-value threshold) in the 4th independent quarter of the discovery population. PRS was also performed using only clumped variants passing genome-wide significance.

These clumped GWS SNPs were also used to create trans-ethnic PRS models. As expected (Martin et al. 2019; Amariuta et al. 2020), cross-population PRS prediction performed worse than inter-population prediction. SNPs were filtered based on Minor Allele Frequency and imputation INFO score in the two populations. To further improve cross ancestry prediction, SNPs in LD with lead SNPs (proxies) were also considered for prediction in multiple frameworks, first including proxies defined using LD calculated in the discovery ancestry and then using proxies defined using the prediction ancestry, as shown in Supp Figure 4. Prediction from UKB to CKB improved beyond inter-CKB prediction; we believe this is likely due to the larger discovery sample size of UKB. However, using proxy information failed to improve prediction accuracy for trans-ethnic PRS both when discovering in UKB and predicting in CKB as well as when discovering in CKB and predicting in UKB.

## Data Availability

The individual-level genotype and phenotype data are available from the UKBB by application at https://www.ukbiobank.ac.uk/. GWAS summary statistics will be available from the GWAS catalog https://www.ebi.ac.uk/gwas/. CKB baseline data are available under the CKB Open Access Data Policy, full details at www.ckbiobank.org. Sharing of CKB genotyping data is currently constrained by the Administrative Regulations on Human Genetic Resources of the People’s Republic of China. Access to these and certain other data is available through collaboration with CKB researchers.

## Acknowledgements

This research was conducted using the UK Biobank Resource under application 31063. We thank participants of the UK Biobank for making this work possible. This work was funded in part by NIH awards T32 HG002295 and R01DK075787. The CKB baseline survey and the first re-survey were supported by the Kadoorie Charitable Foundation in Hong Kong. Long-term follow-up was supported by the Wellcome Trust (212946/Z/18/Z, 202922/Z/16/Z, 104085/Z/14/Z, 088158/Z/09/Z), the National Natural Science Foundation of China (82192901, 82192904, 82192900), and the National Key Research and Development Program of China (2016YFC0900500). DNA extraction and genotyping was funded by GlaxoSmithKline and the UK Medical Research Council (MC-PC-13049, MC-PC-14135). The project is supported by core funding from the UK Medical Research Council (MC_UU_00017/1,MC_UU_12026/2, MC_U137686851), Cancer Research UK (C16077/A29186; C500/A16896), and the British Heart Foundation (CH/1996001/9454) to the Clinical Trial Service Unit and Epidemiological Studies Unit and to the MRC Population Health Research Unit at Oxford University. We thank the participants in CKB and the members of the survey teams in each of the 10 regional centres, and to the project development and management teams based at Beijing, Oxford and the 10 regional centres. China’s National Health Insurance provides electronic linkage to all hospital treatments. The content is the sole responsibility of the authors and does not necessarily reflect the official views of the NIH.

## Supplementary figures: full list

1. Lambda GC for GWAS

2. Collider bias

3. LDSC heritability for each trait in 2 populations and with sample size match

4. GSEA of each phenotype

5. HOX gene significance

6. FM concordance when taking new approach

7. FM statistics

8. Height SNPs: SHR p-values nearby (and SH vs Leg, and vise versa)

9. Yengo 2022 effects

10. Linkage for SHR signals to nearby height signals

11. Leg/SH effects of classified Height CS SNPs

12. SNP classification gene set proximities for pheno pairs

13. Fine-mapping-overlapping gene growth plate expression specificity

14. Specific loci examples of phenotype FM overlap

15. Ancestry phenotype differences

16. WCC efficacy

17. WCC correction with effect size comparisons

18. Ancestrally heterogeneous SNP effect sizes

19. Specific examples of ancestry FM heterogeneity

20. Schematic, PRS, PRS xAncestry

21. Table 1: FM summary

22. Table 2: MA results x 4 phenotypes

23. Table 3: Gene prioritization

24. Table 4: Gene set prioritization

25. Table 5: Gene classifications

26. Table 6: FM summary statistics x 4 phenotypes

---

**Supplemental figure 1:**
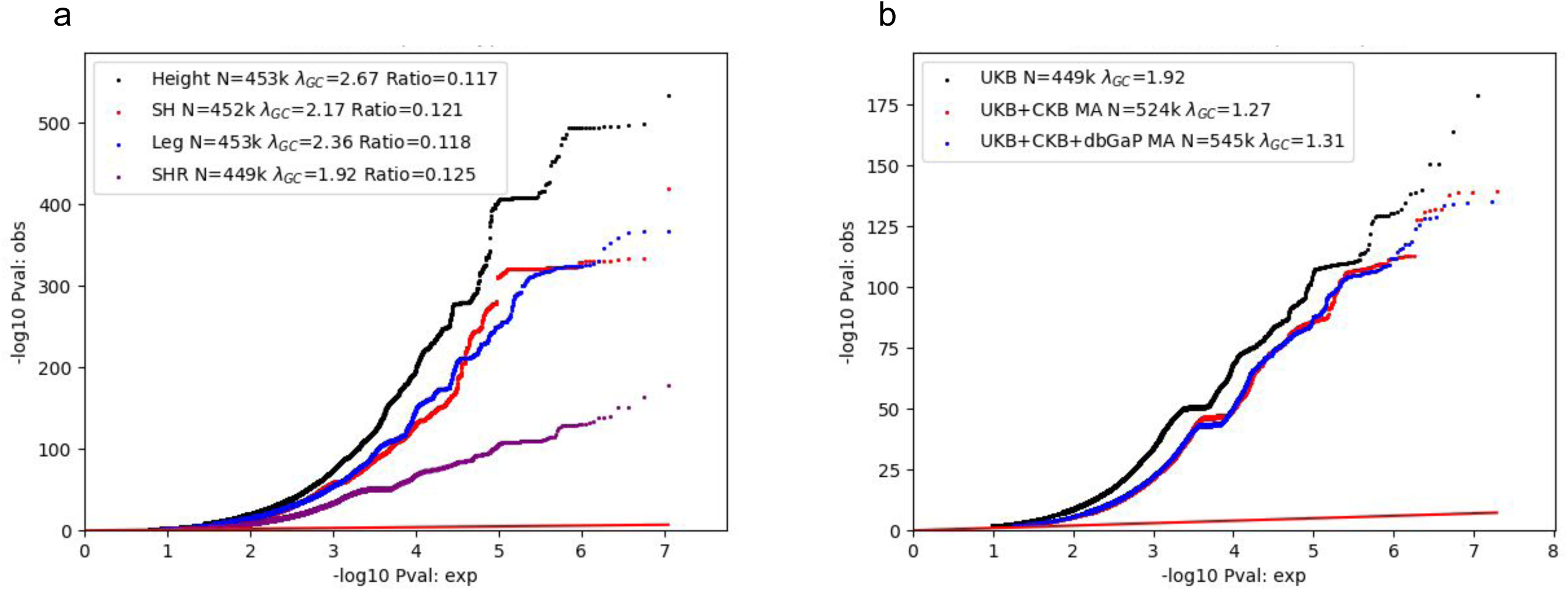
A. QQ for GWAS associations performed in UKB for phenotypes. Lambda GC values observed above 1; LDSC Ratio values are within normal bounds and are less than 0.2. B. Q for SHR GWAS associations in samples UKB, UKB meta-analyzed with CKB, and UKB and CKB meta-analyzed with results from Chan 2015.

**Supplemental figure 2:**
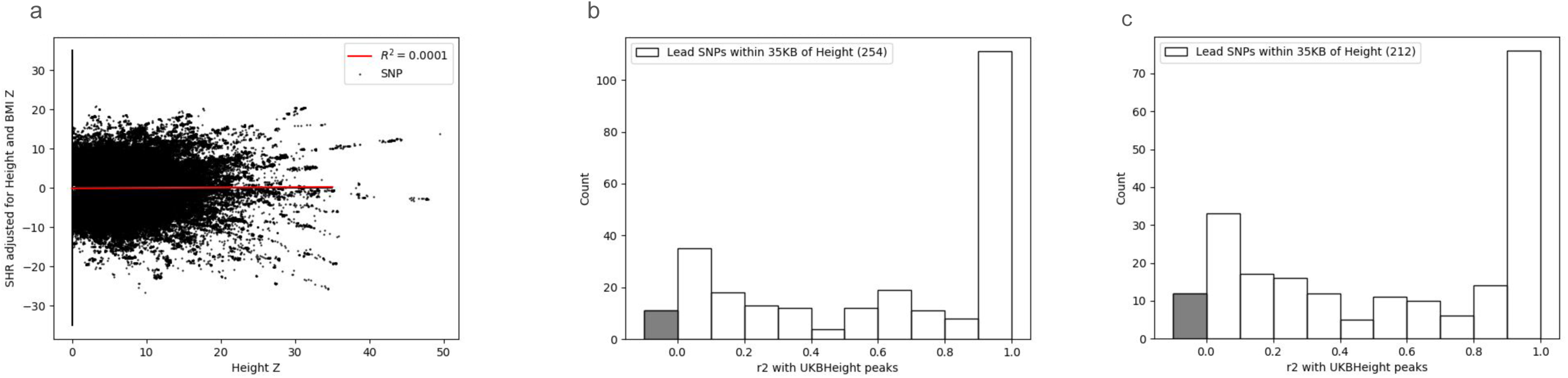
A. Z score comparison for SNPs associated with height (x-axis) and SHR (y-axis) in UKB. R2 = 0.01. Red line marks line of best fit. B-C. LD (R2) from SHR (B) and SHR-unadjusted (C) lead SNPs to height lead SNPs, limited to SHR or SHR-unadjusted SNPs within 35Kbp of a height lead SNP. Grey bar marks SNPs not near height SNPs (LD < 0.01). LD calculated using 1000 Genomes.

**Supplemental Figure 3:**
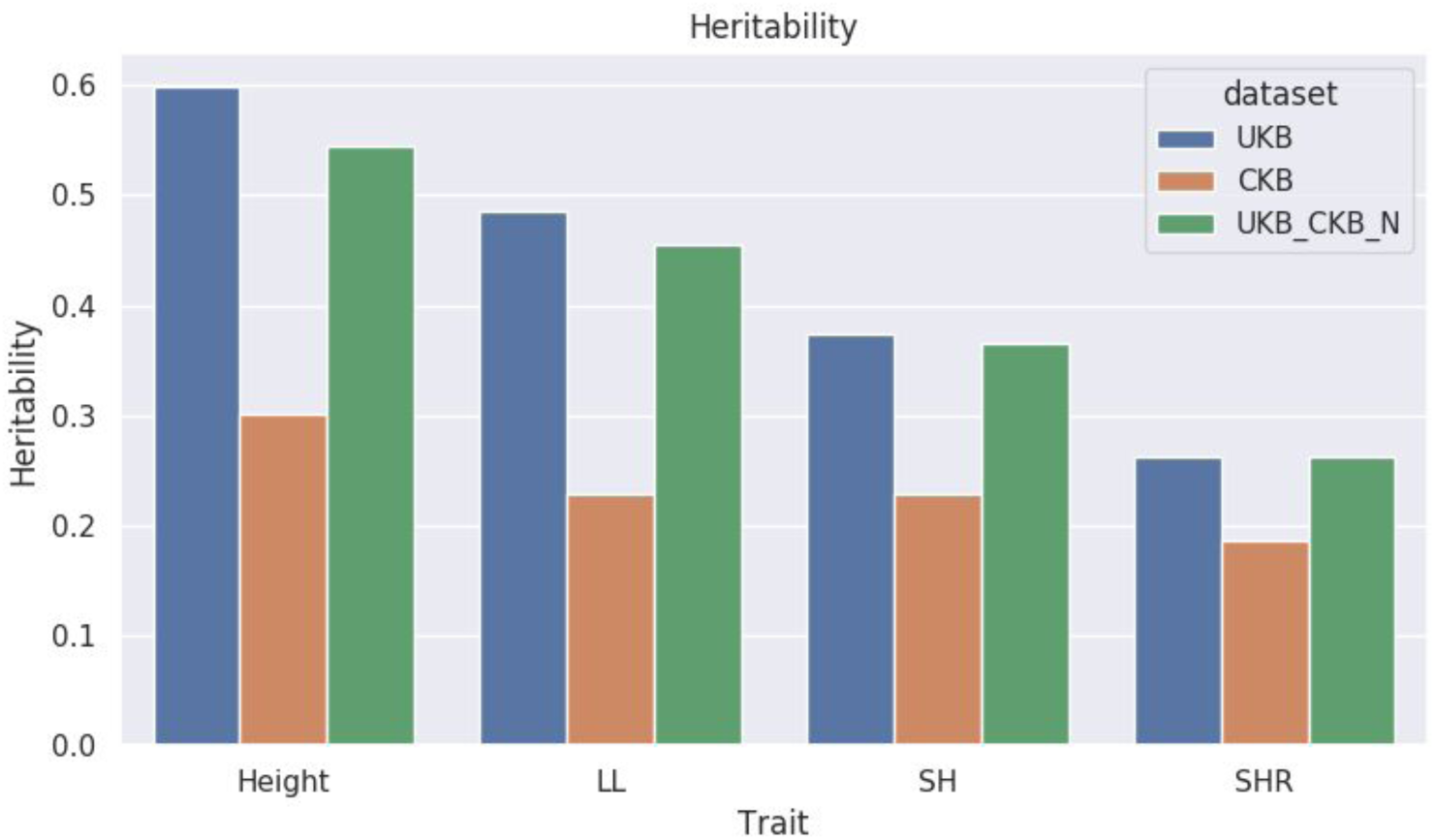
Heritability estimates for height-related phenotypes in samples UKB, CKB, and UKB down-sampled to match CKB. Heritability calculated using LDSC.

**Supplemental figure 4:**
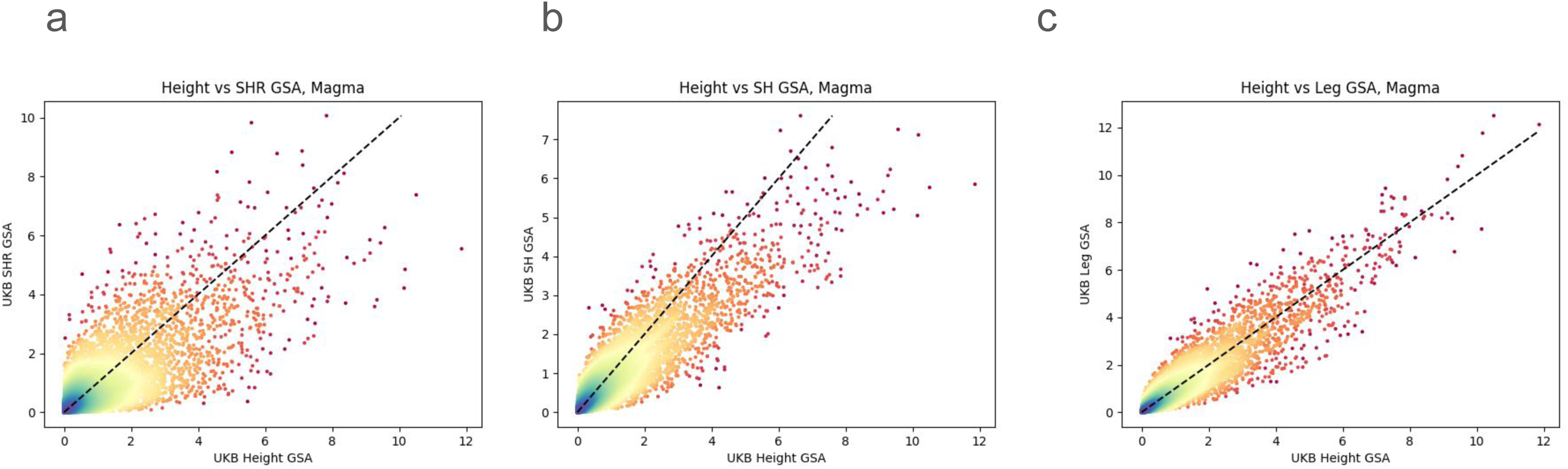
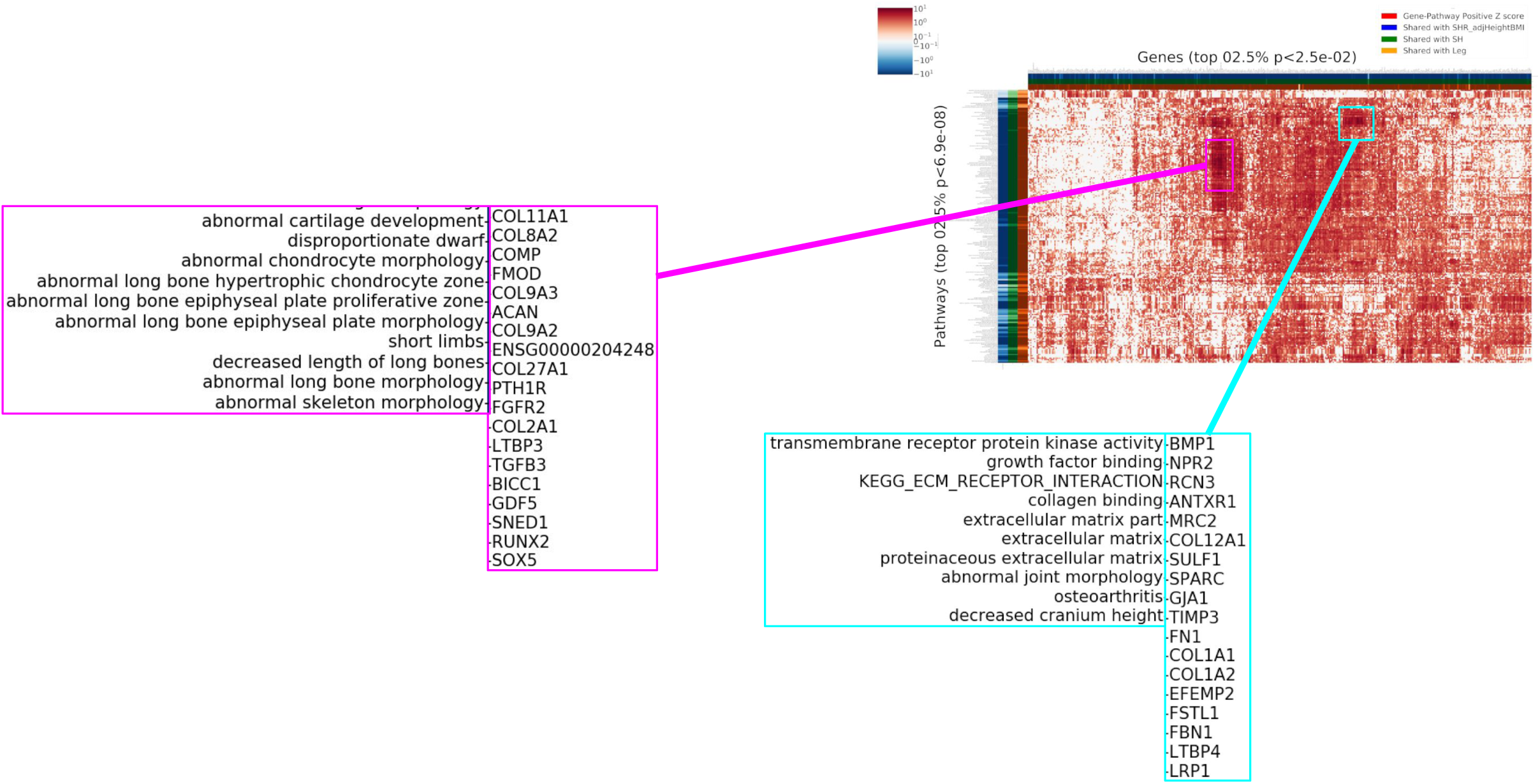
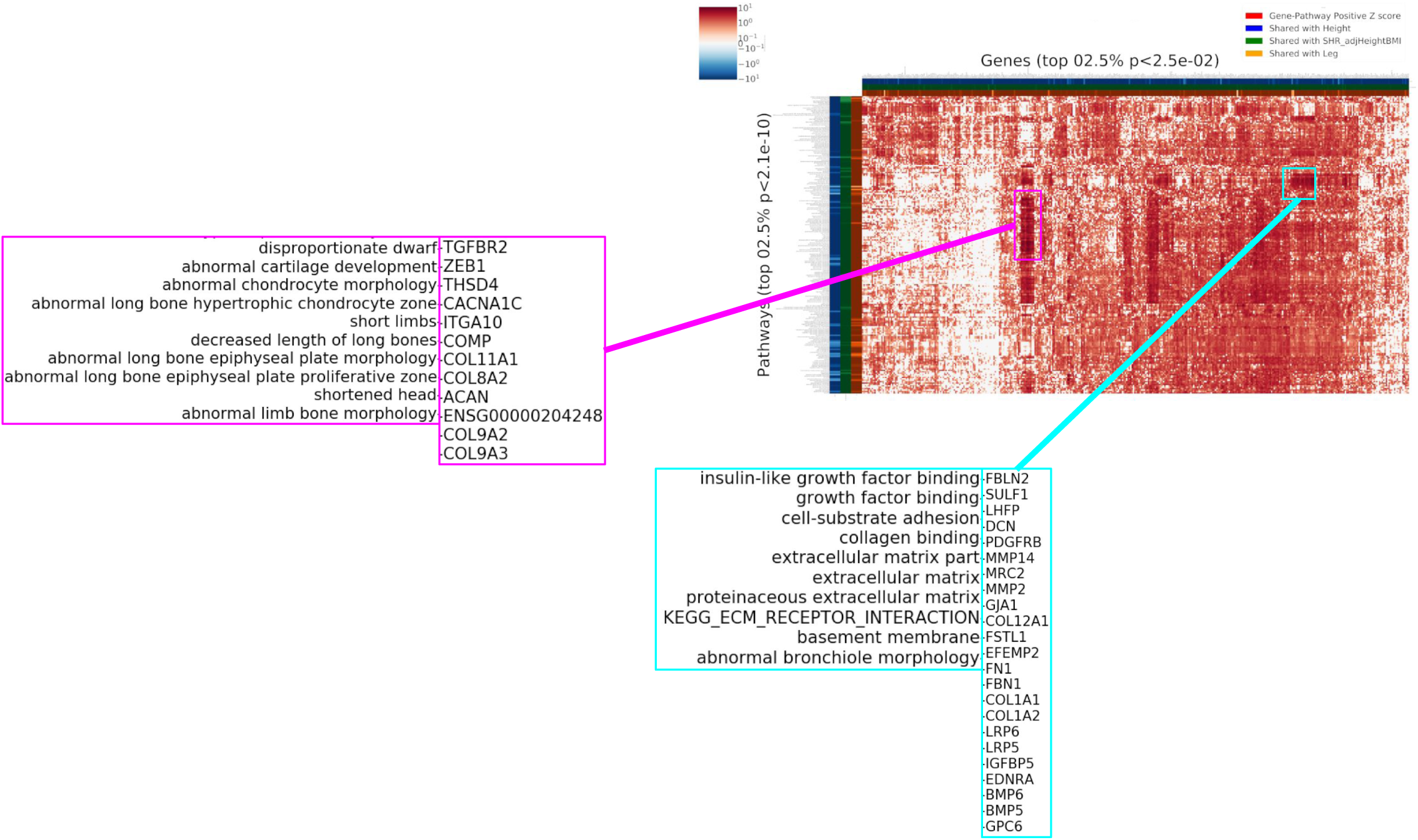
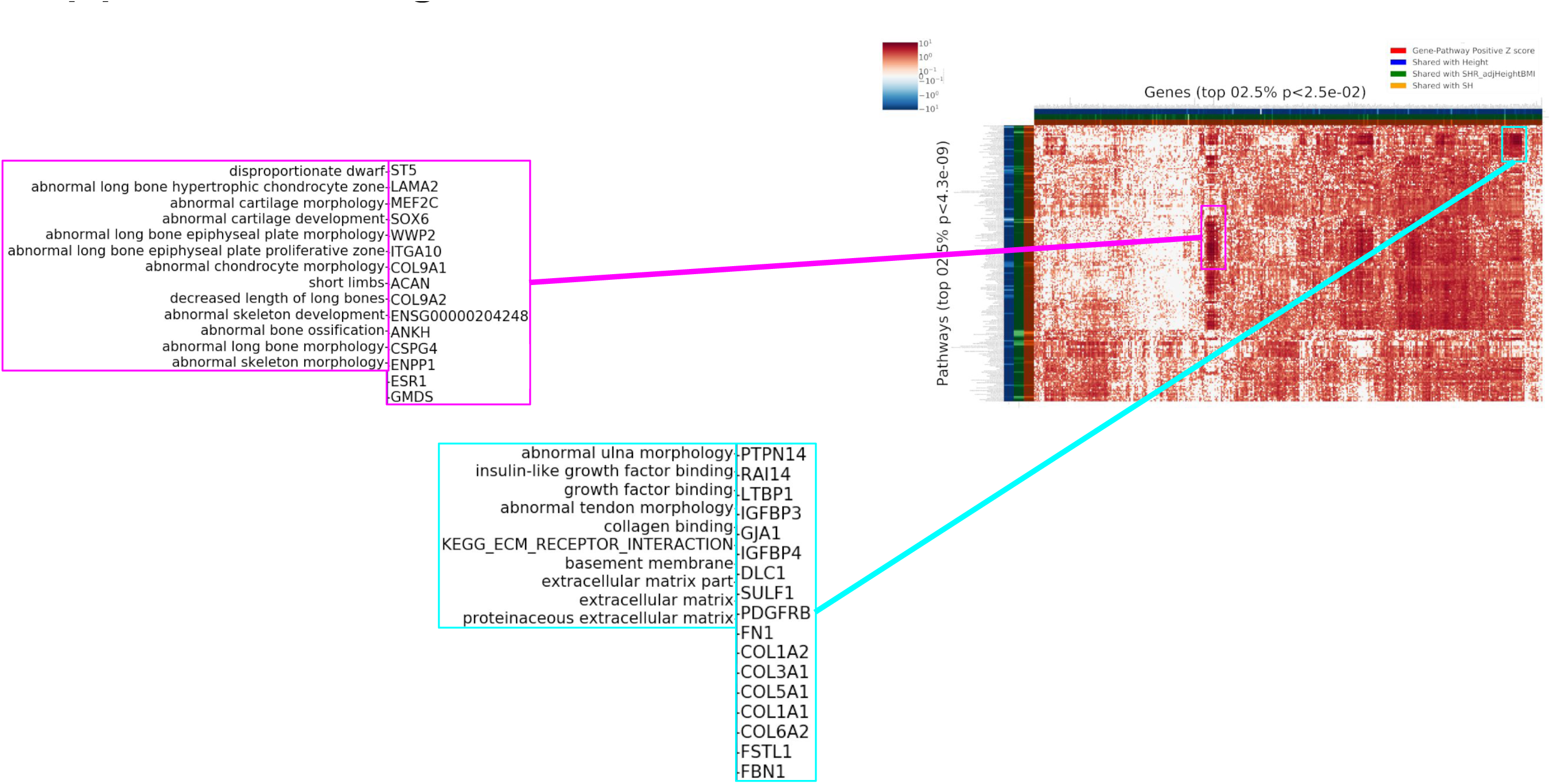
GSEA results for four phenotypes. A,B,C. Gene-set significance for height (x-axis) compared with SHR, SH, and LL respectively (y-axis). Color indicates point density. D,E,F. Each row is a gene set identified by enrichment analysis (GSEA) of height, SH, and LL, respectively, using MAGMA (top 2.5% of prioritized gene sets), and each column is a gene prioritized by PoPs (top 2.5% of prioritized gene sets). The squares in the heatmap indicate the likelihood of membership of the gene in the corresponding gene set (gene-pathway Z-score supplied by the DEPICT reconstituted gene sets Pers 2015). Row and column annotations in the left and top margins indicate whether the gene set or gene, respectively, was also prioritized prioritization by similar analysis of each of the other height-related phenotypes (colored indicates prioritized). Zoomed regions show clusters of related prioritized genes and gene sets.

**Supplemental figure 5:**
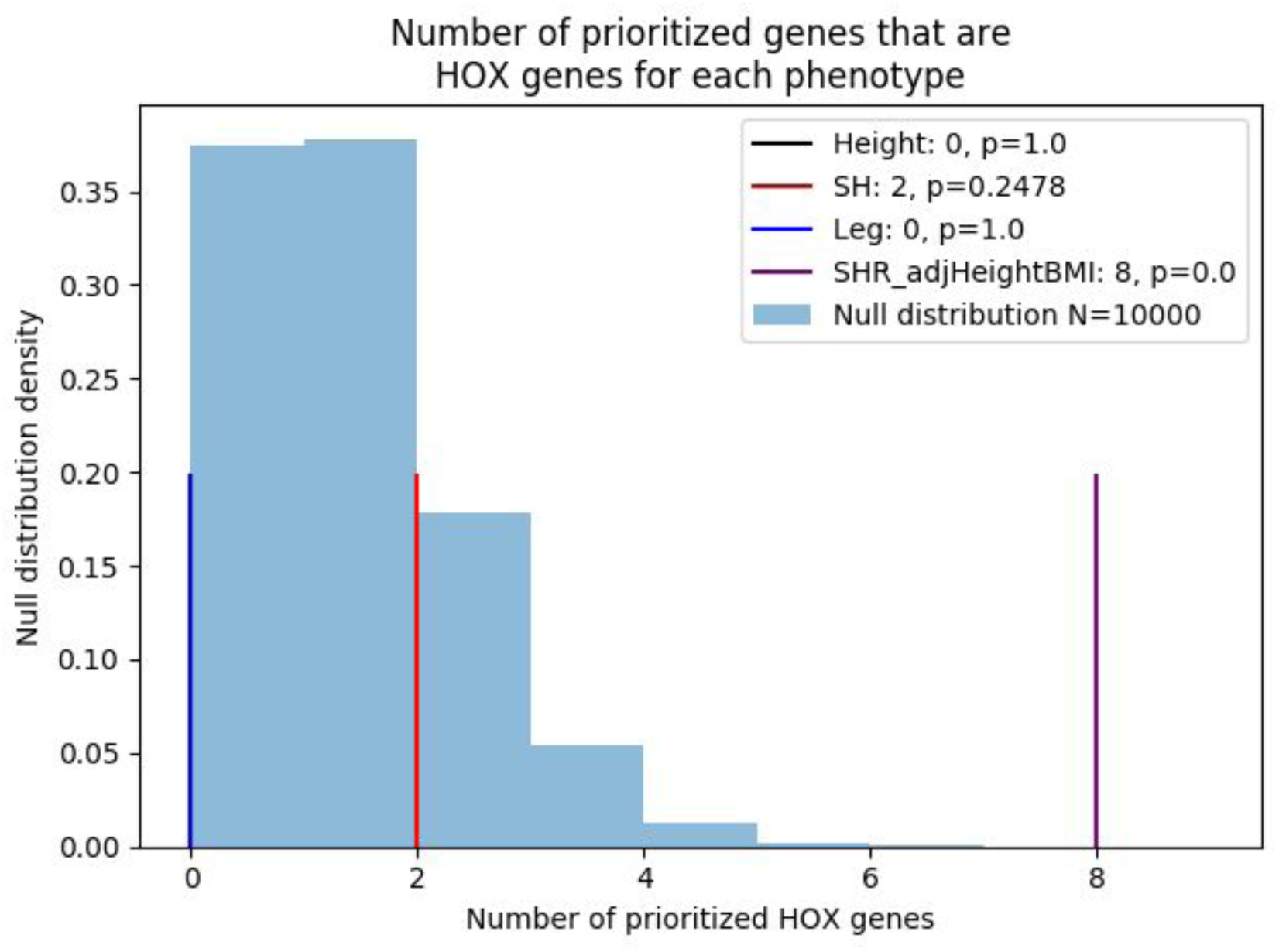
SHR (purple observation) has more HOX genes than the other height-related phenotypes, and significantly more than random prioritization.

**Supplemental figure 6:**
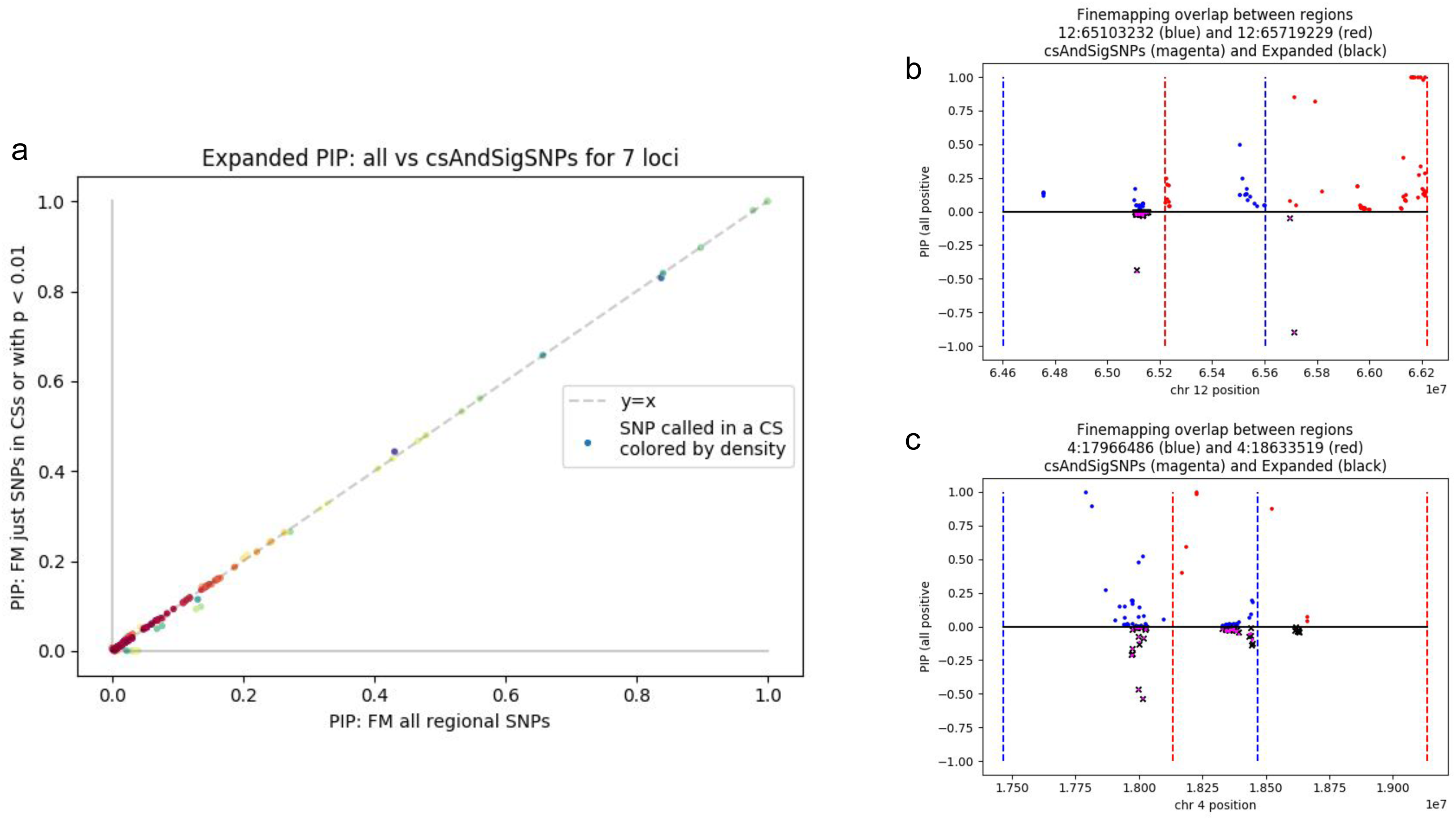
A. Consistency of fine-mapping of regions greater than 1MB when including all (MAF and info filtered) SNPs in SuSIE model (x-axis) vs including only a filtered set of SNPs (in a CS in 1MB analysis, OR p-value < 0.01) (y-axis). B, C. Comparison of fine-mapping results for regions showing consistency and lack of consistency, respectively, between a complete SuSIE model (all SNPs) and a filtered SuSIE model (only SNPs in CSs or passing a p-value filter). Top, blue and red: 1MB window fine-mapping of overlapping regions. Bottom, black: complete SuSIE model. Bottom, pink: filtered SuSIE model. PIP values are positive but shown as a Miami plot for ease of comparison.

**Supplemental figure 7:**
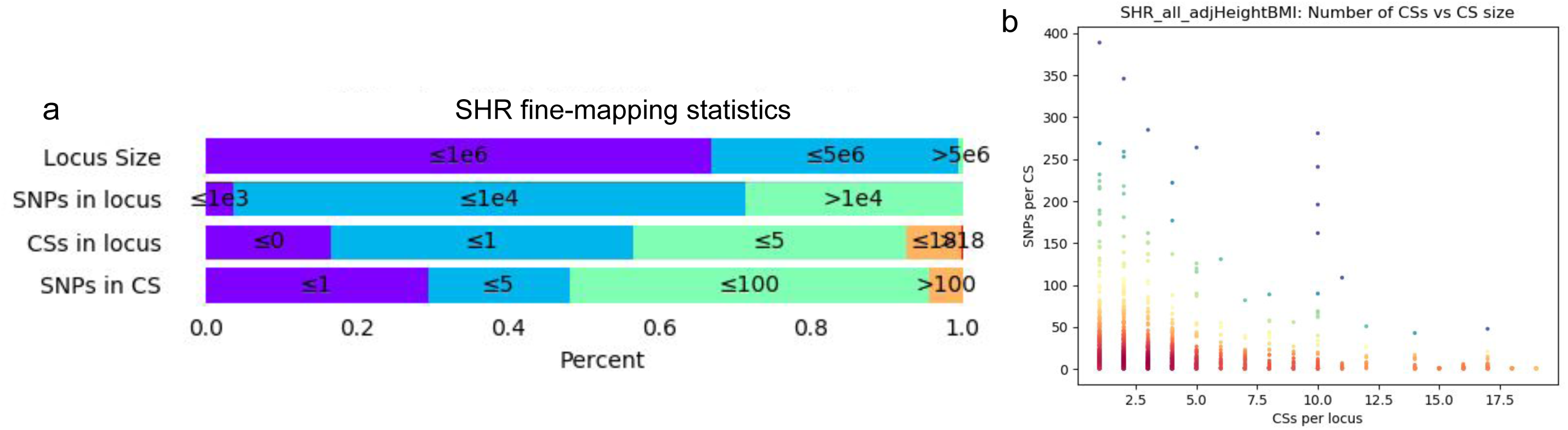
A. Finemapping was performed for traits height, SHR, SH, and leg. For SHR, approximately ⅔ of loci were evaluated as 1MB, with the remaining loci resulting from combining overlapping finemapped regions containing genome-wide-significant associations. For the majority of loci, at least 1000 SNPs were evaluated. Approximately 30% of loci had 1 CS, and almost all loci had 18 or fewer CSs. About half of CSs had 5 or fewer SNPs, with a minority of CSs containing many SNPs (above 100). B. Most CSs have few SNPs, and most loci few CSs. as expected by this overlap, many loci have few CSs, each with few SNPs.

**Supplemental figure 8:**
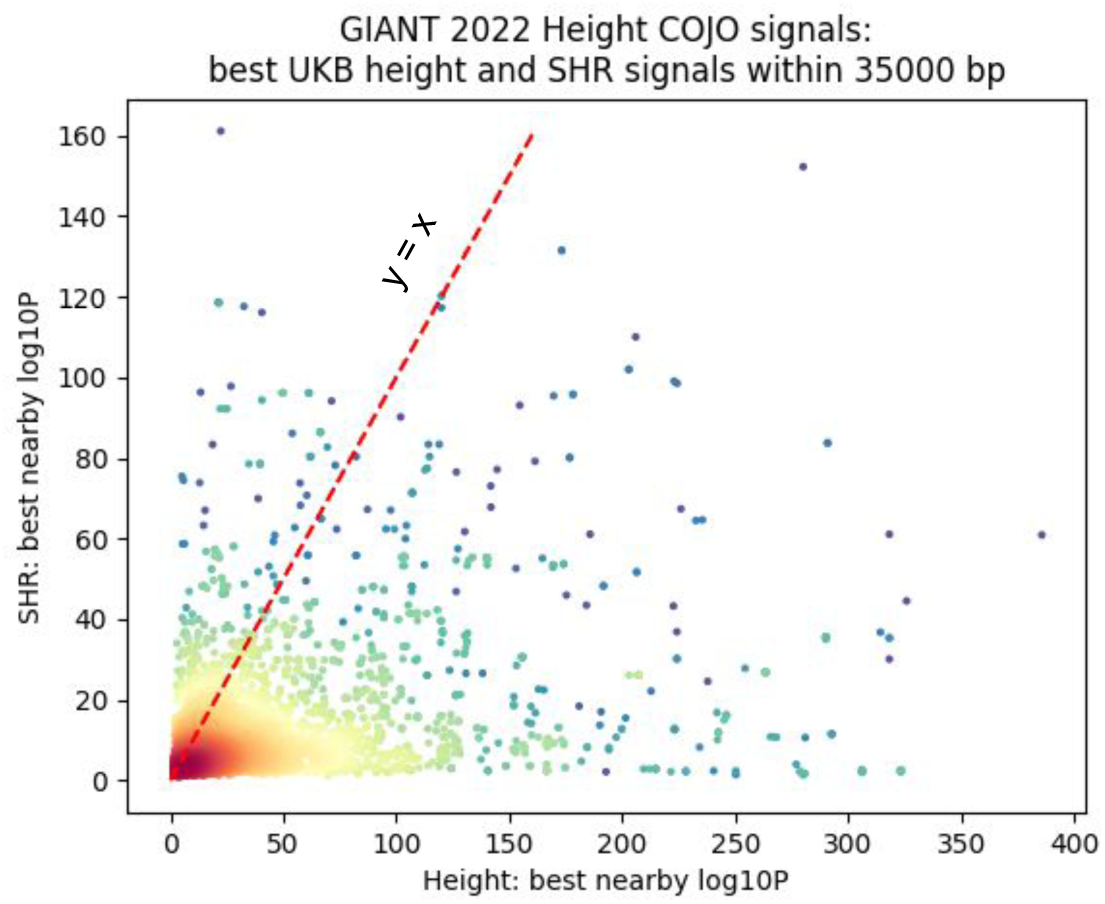
A. For each of 12111 COJO signals identified in Yengo 2022, most significant nearby p-values from height and SHR GWAS were identified. The y=x line is marked in red. Most SNPs have more nearby significance for height than SHR, but there exist also many SNPs containing more nearby significance for SHR.

**Supplemental figure 9:**
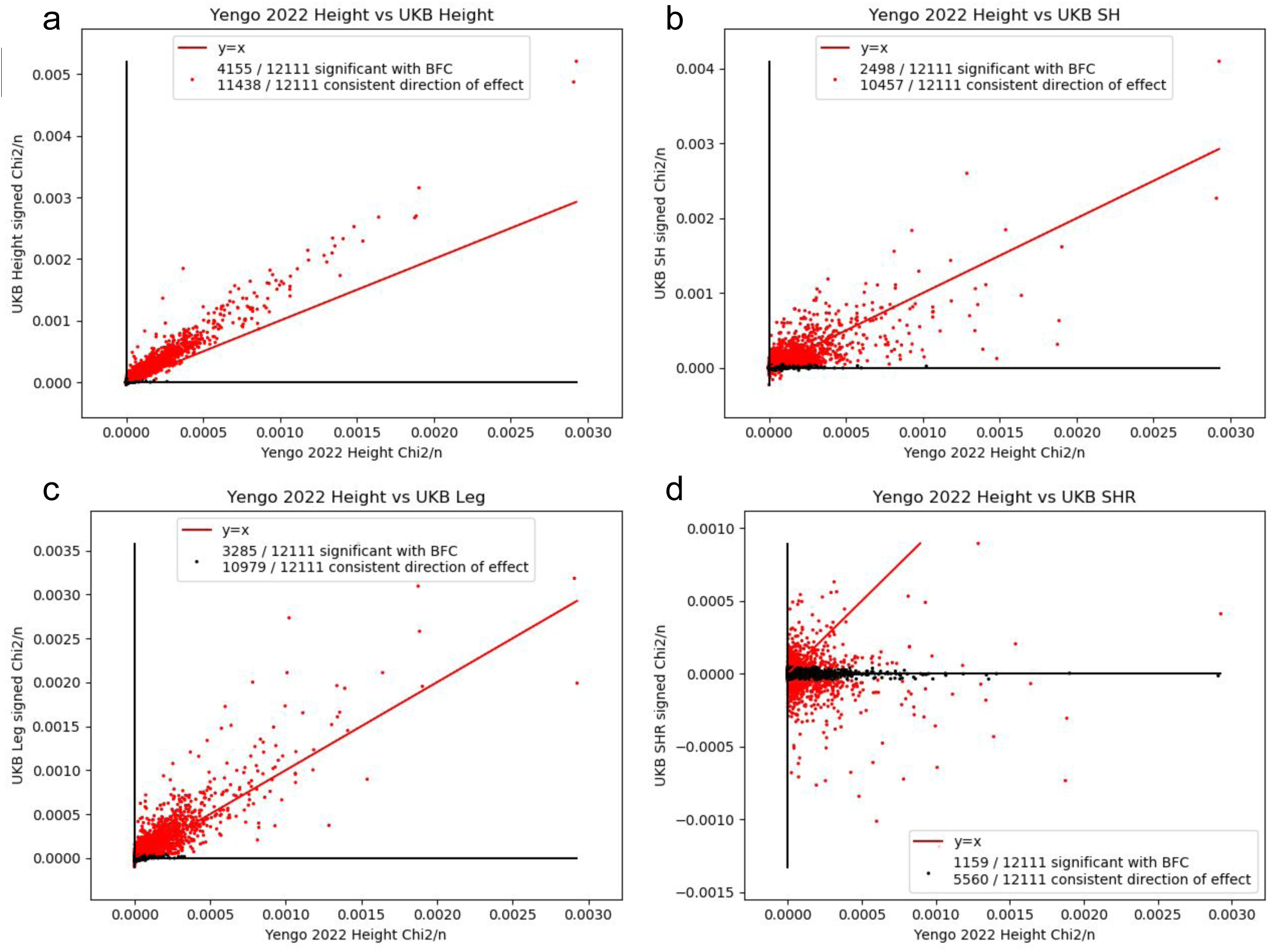
A,B,C,D. For each of 12111 COJO signals identified in Yengo 2022, the estimated effect size in GWAS performed in UKB for height and related phenotypes SH, LL, and SHR respectively. SNPs in red are significantly associated with the UKB phenotype after bonferroni correction for number of SNPs.

**Supplemental figure 10:**
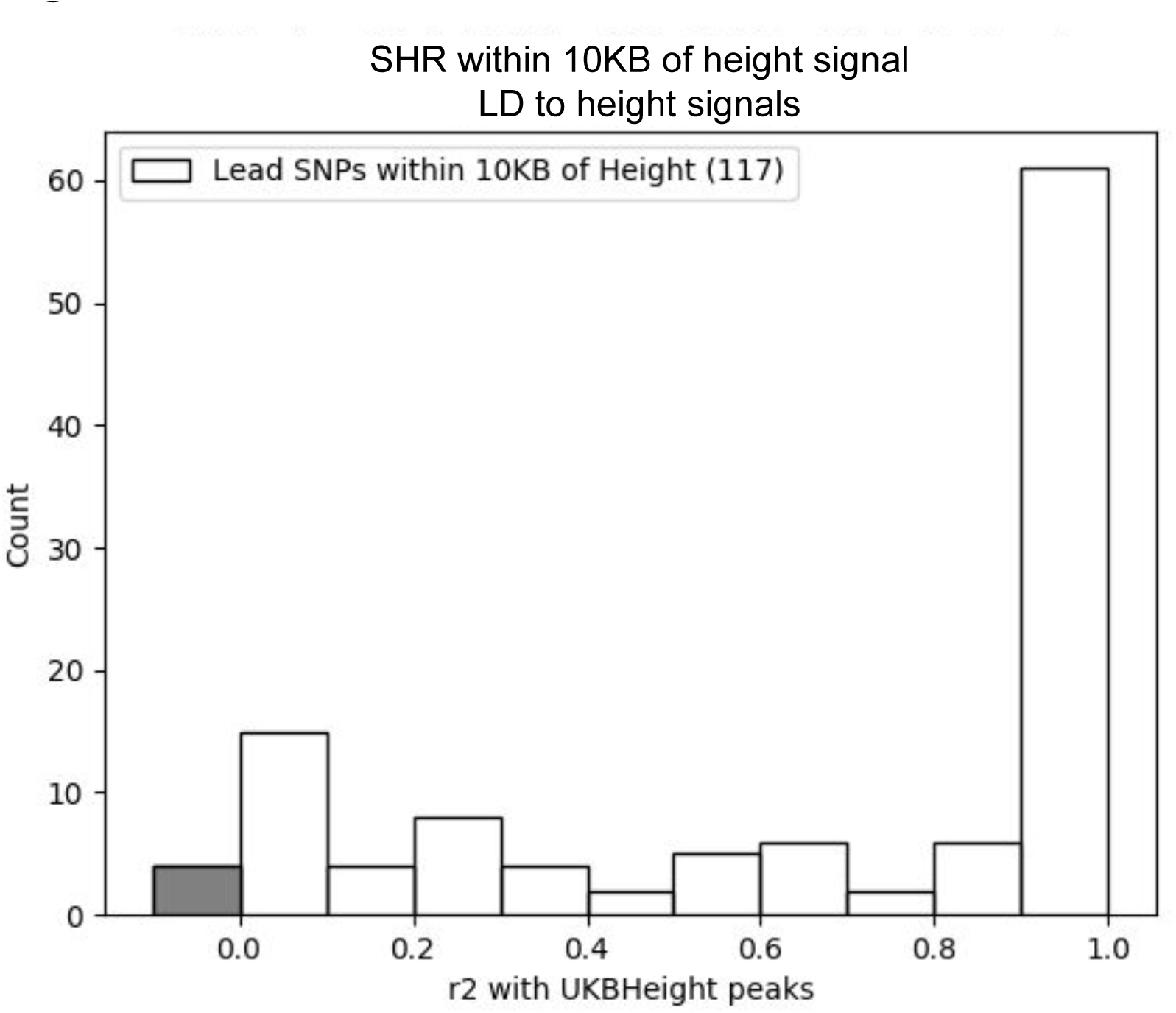
SHR lead SNPs were filtered for those near (within 10kb) of a UKB height lead SNP. LD was then calculated using 1kg EUR between height and SHR leads, and their distribution plotted. SHR leads not in the LD panel are included in the grey bar.

**Supplemental figure 11:**
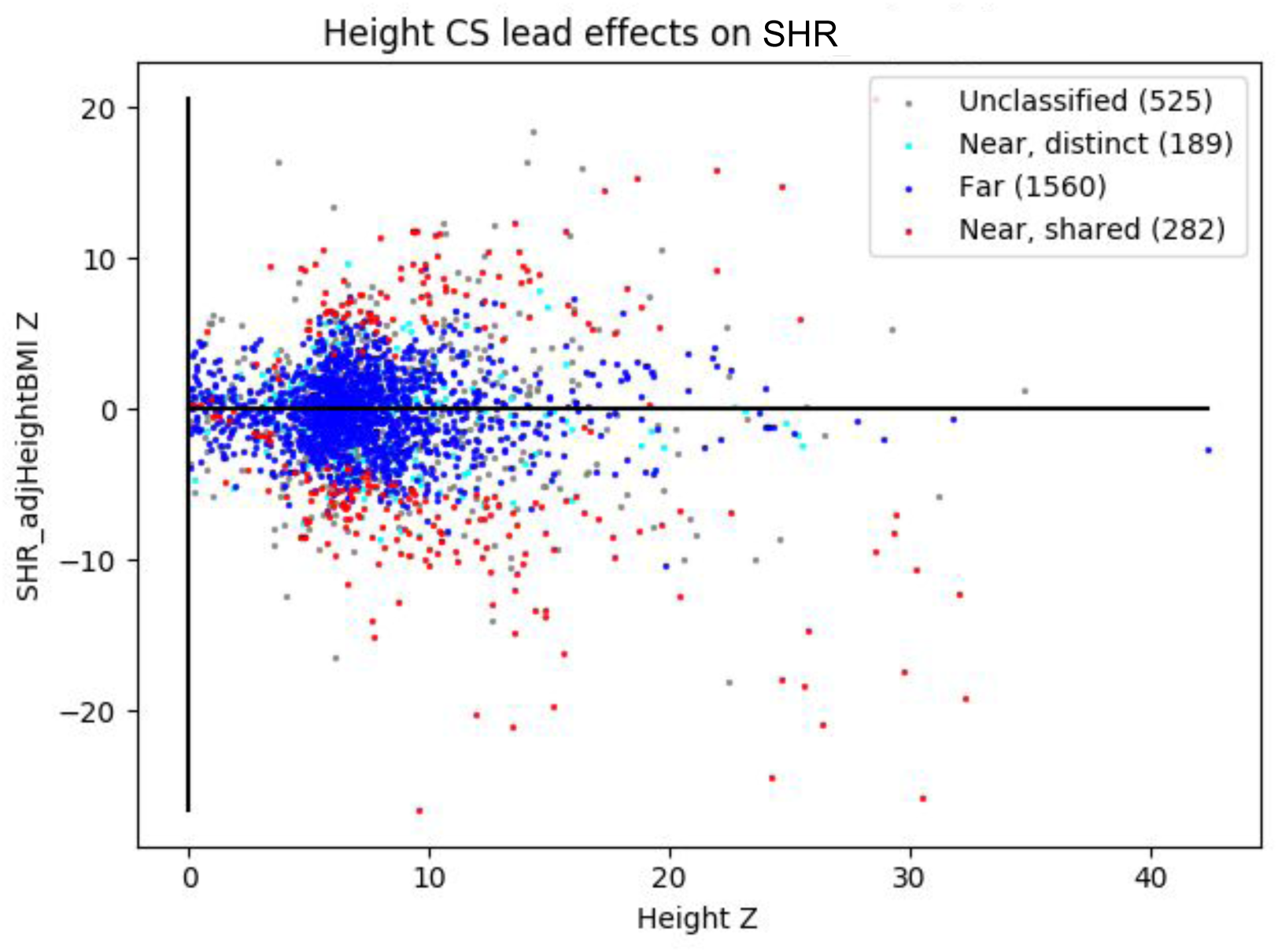
Similar to 2D, for height signals proximal to SHR signals, associations with height and SHR are plotted, with height Z-score on the X-axis and SHR Z-score on the Y-axis. “Near, shared” CSs are plotted in red; “near, distinct” are in black; “far” are in blue.

**Supplemental figure 12:**
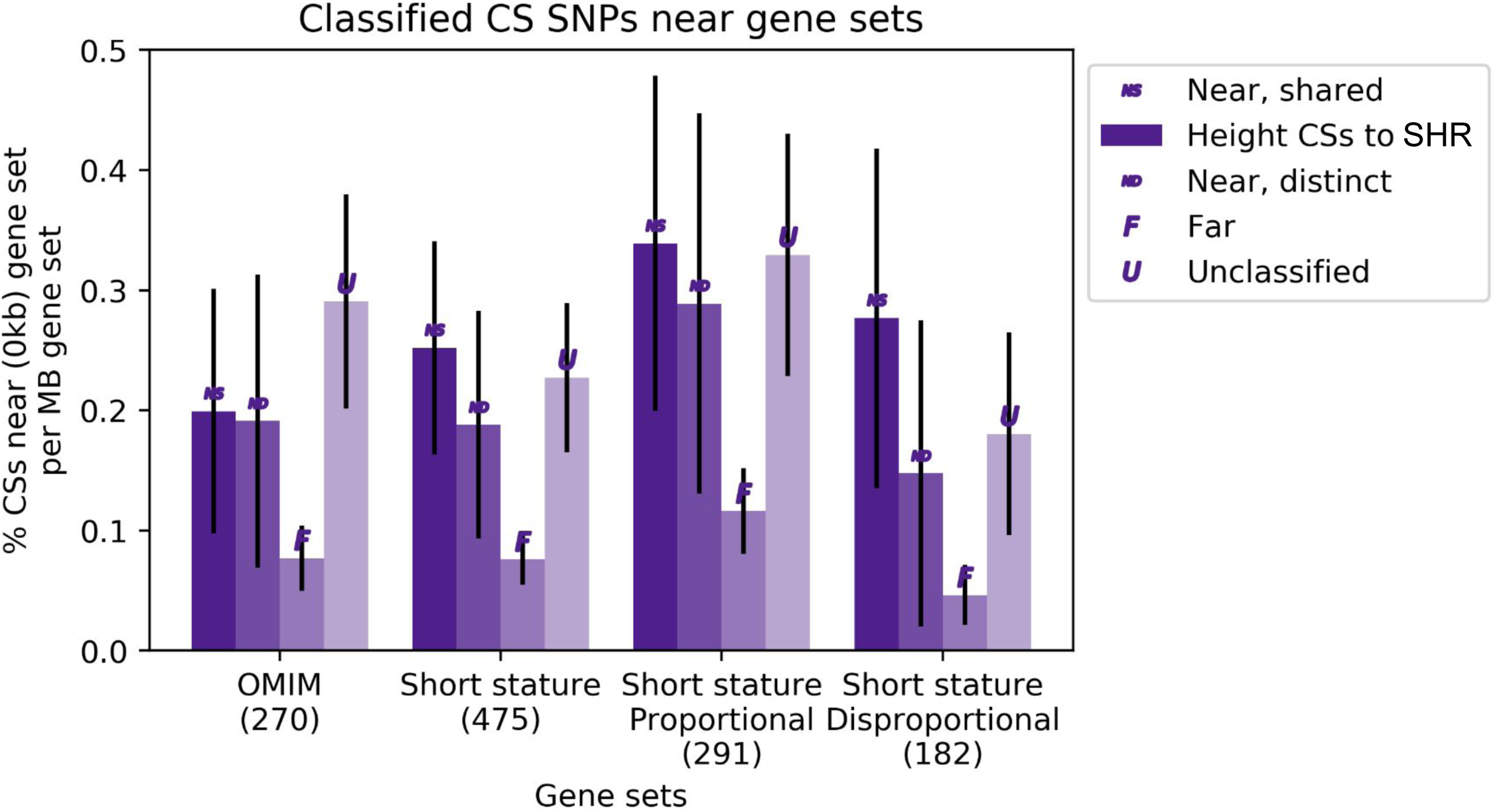
Height CSs’ (classified by their proximity to SHR CSs) overlap with multiple gene sets OMIM, short stature, short stature-proportional, and short stature-disproportional (see results).

**Supplemental figure 13:**
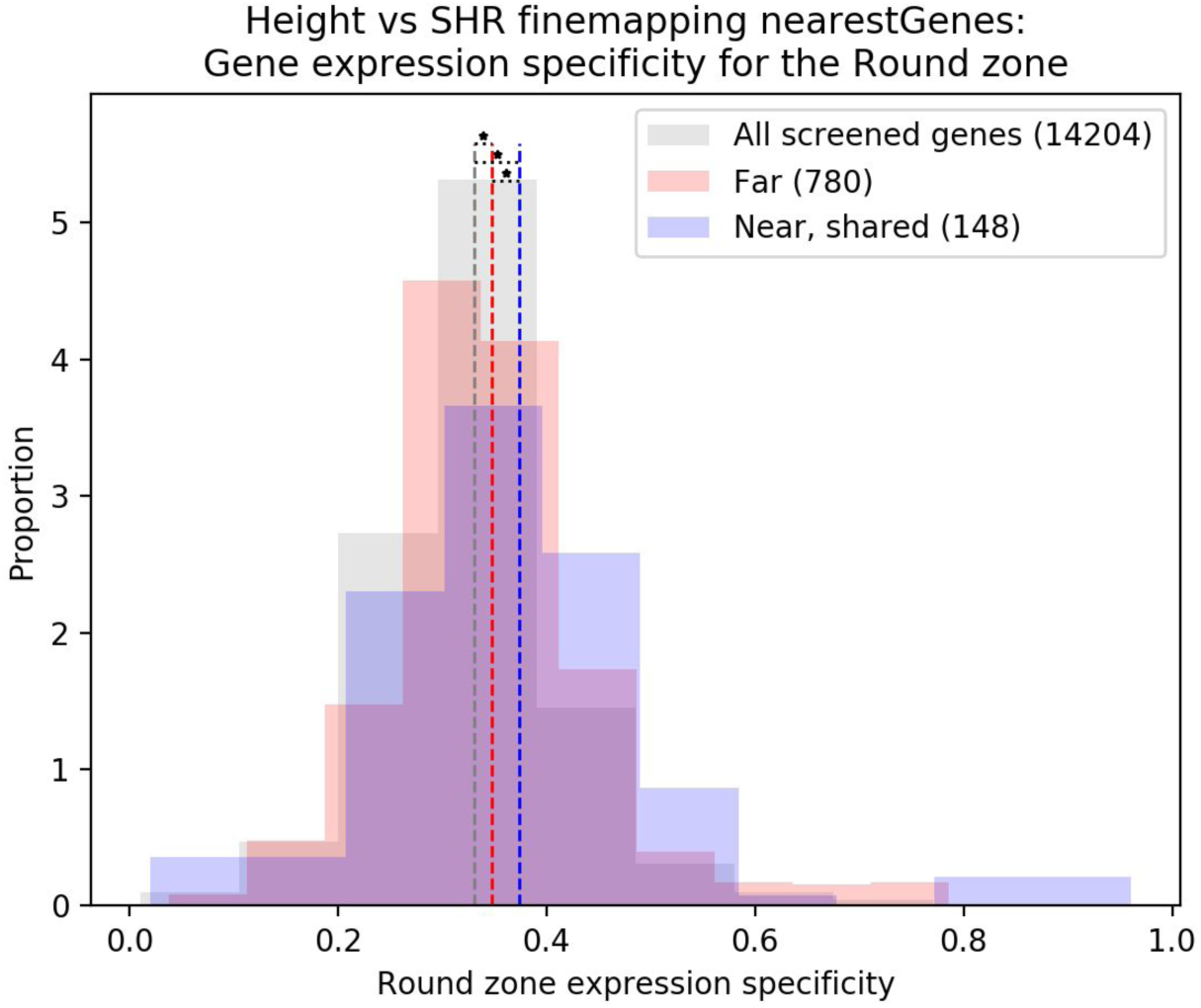
Genes overlapping height CSs’ (classified by their proximity to SHR CSs) are more often specifically expressed in the growth plate’s Round zone rather than Hypertrophic or Flat zones.

**Supplemental figure 14:**
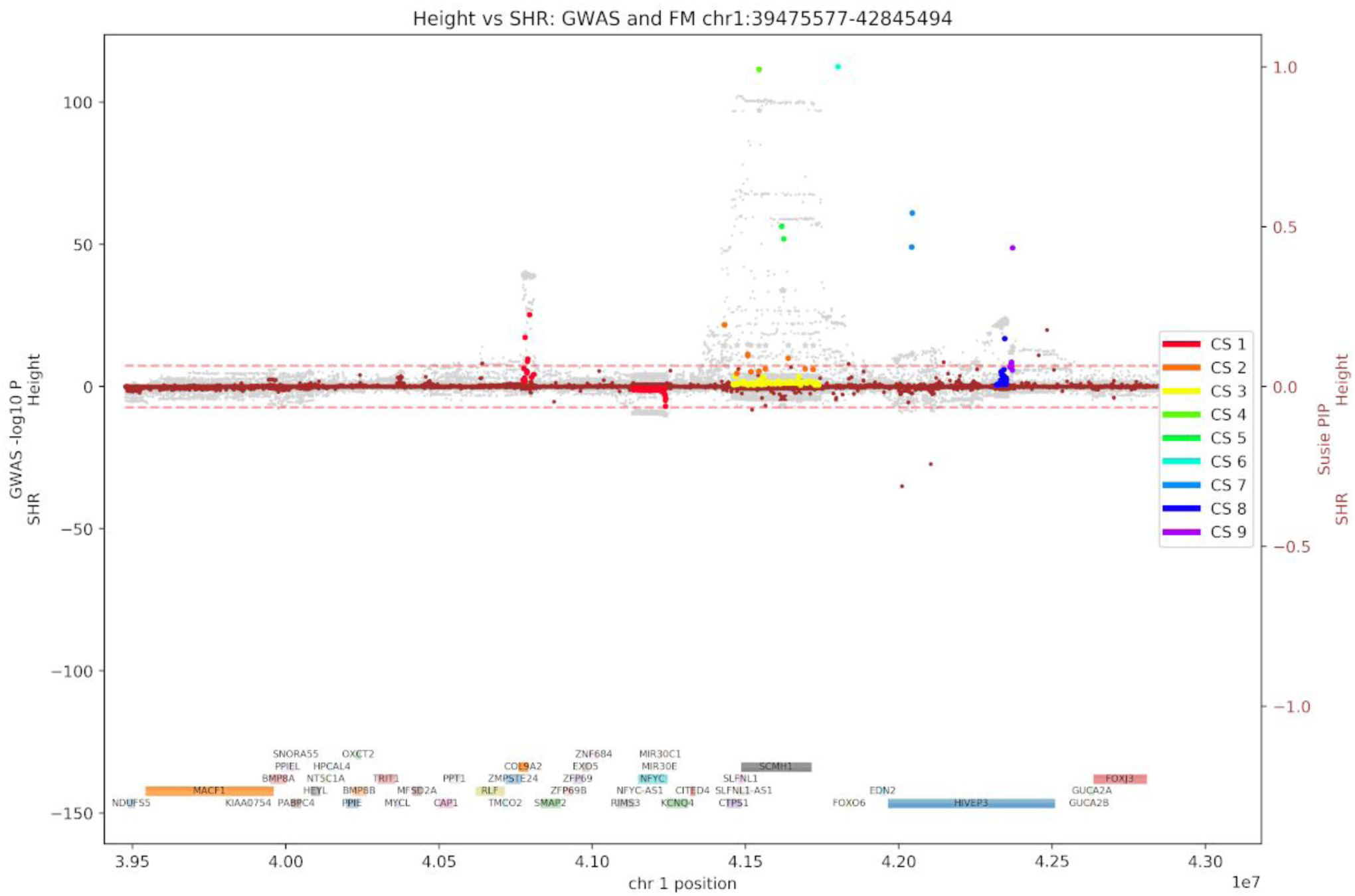
In this example locus on chromosome 1, height GWAS signal is not matched by SHR GWAS signal, and the SHR CSs and height CSs have no counterpart.

**Supplemental figure 15:**
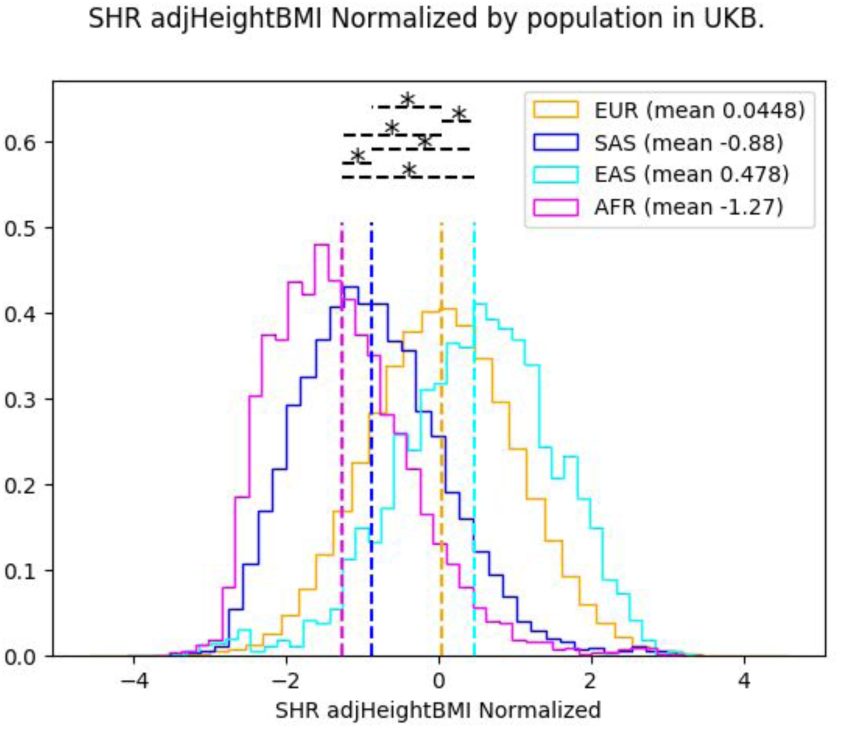
Inverse-normally transformed residuals for SHR, broken down by individuals’ self-identified population.

**Supplemental figure 16:**
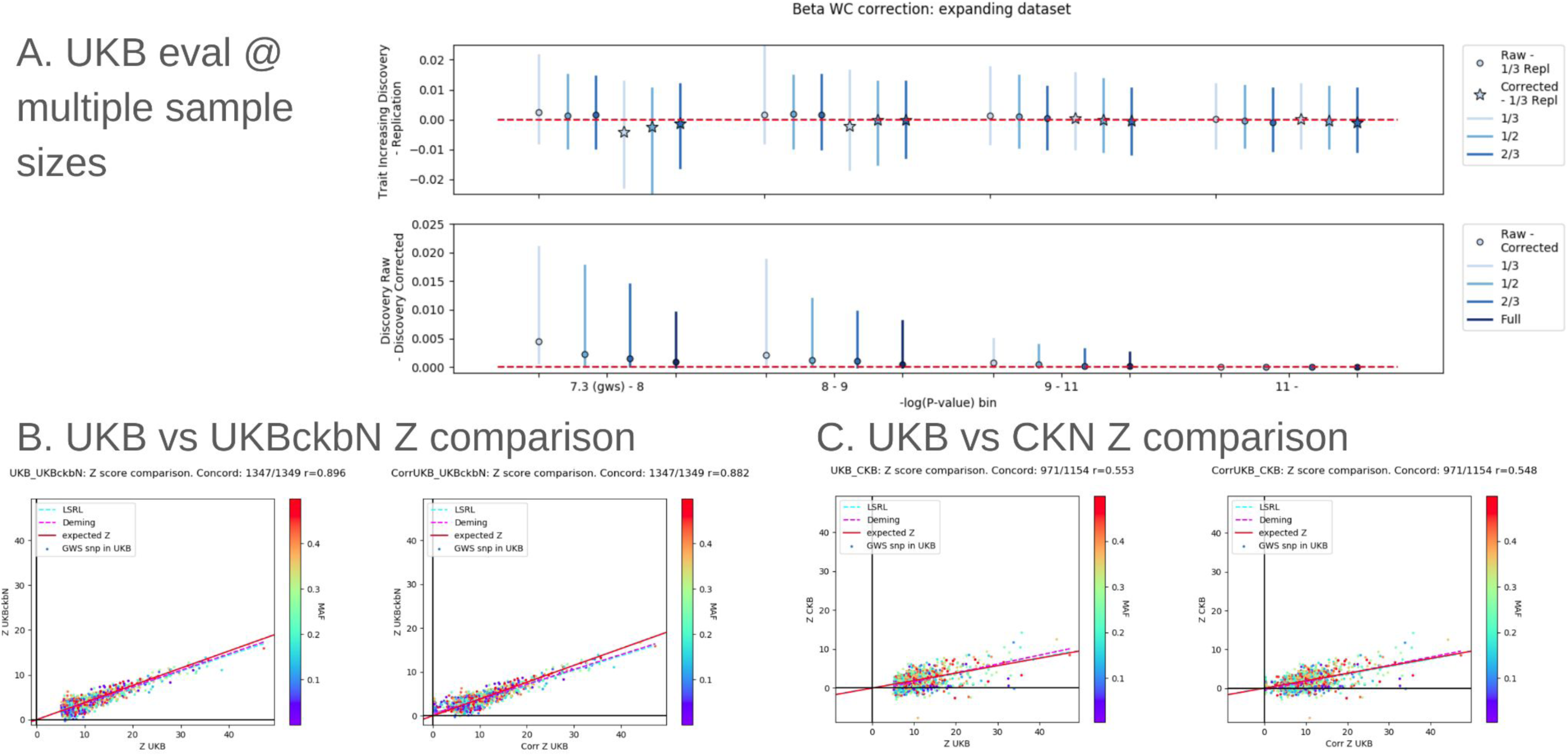
A: Winners curse correction in UKB. Top: Difference between effect sizes in discovery and replication before winners curse correction (circles) and after winners curse correction (stars) across multiple sizes of discovery dataset (blues), broken down by SNP significance (x-axis). Bottom: magnitude of effect size correction across multiple sizes of discovery dataset (blues), broken down by SNP significance (x-axis). B. Z scores at UKB-discovered height lead SNPs estimated in UKB vs a downsampled size-matched UKB subset pre- (left) and post- (right) WC correction. C. Similar to (B), Z scores at UKB-discovered height lead SNPs estimated in UKB vs CKB pre- (left) and post- (right) WC correction.

**Supplemental figure 17:**
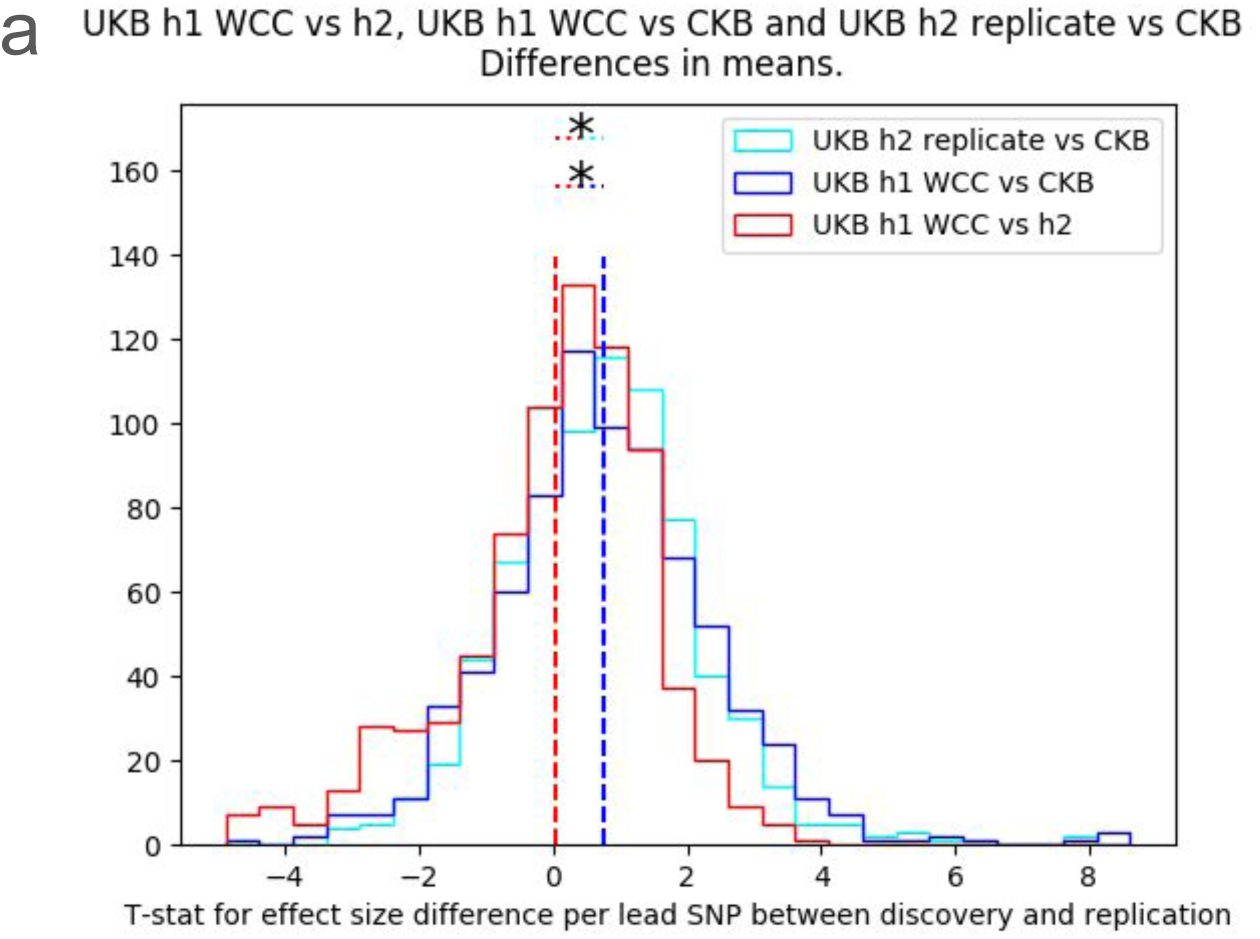
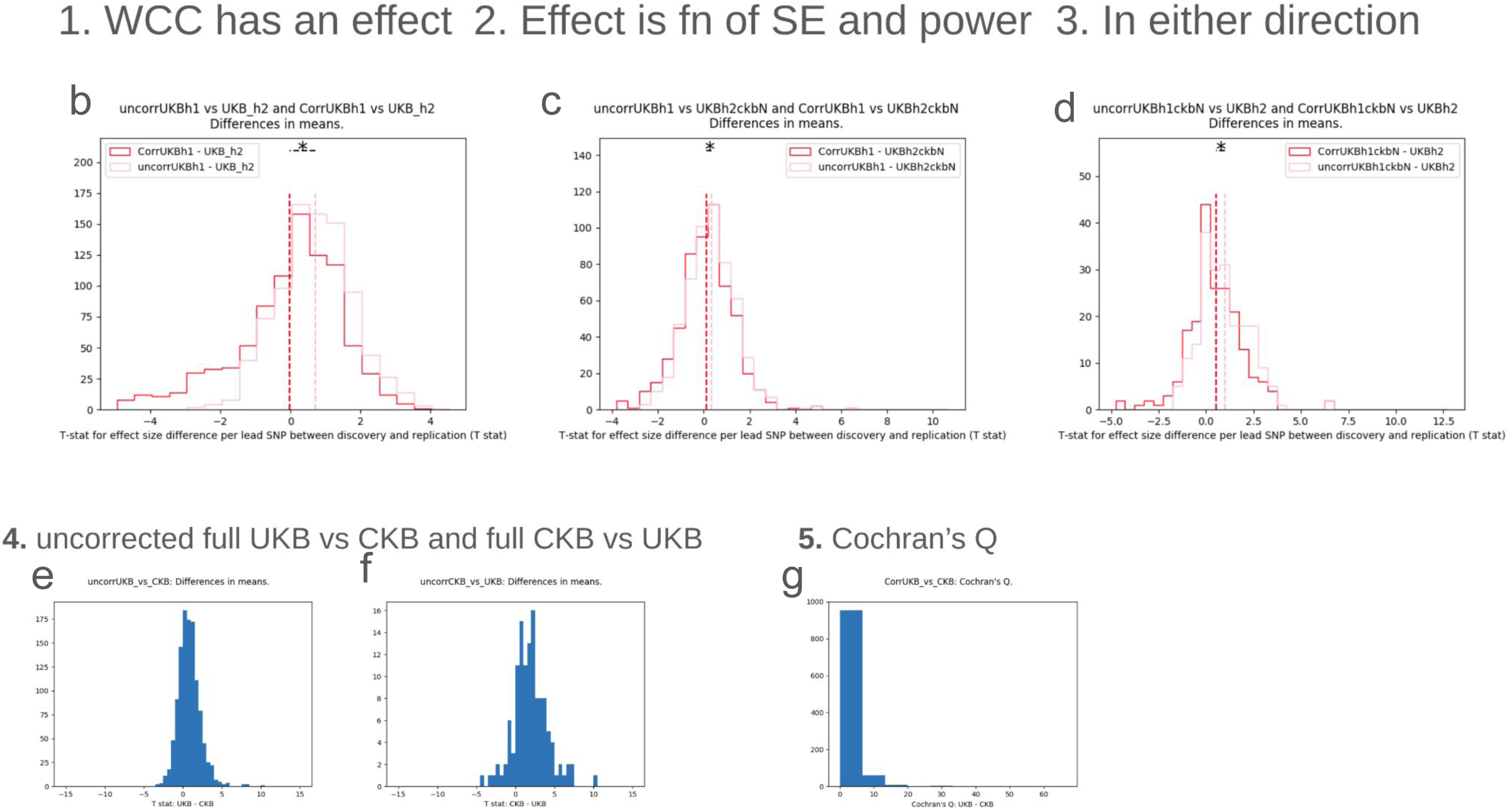
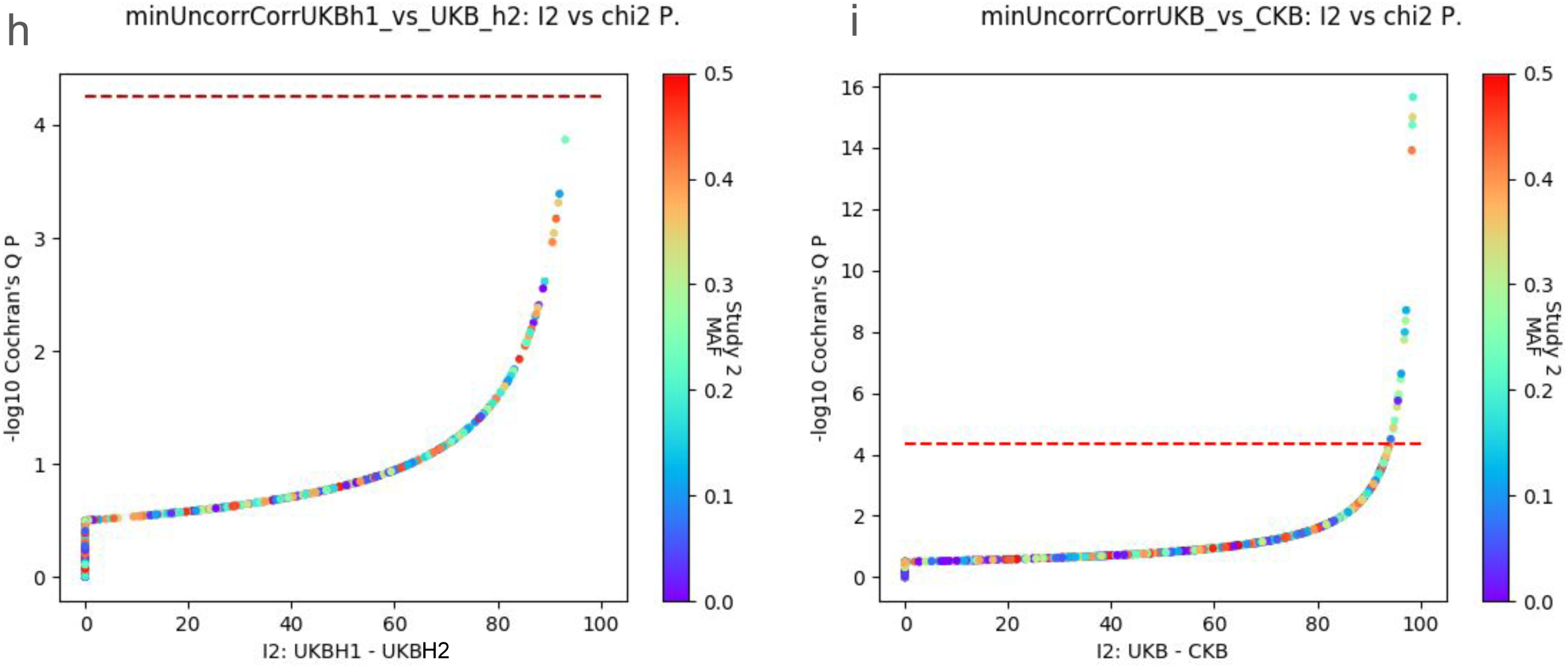
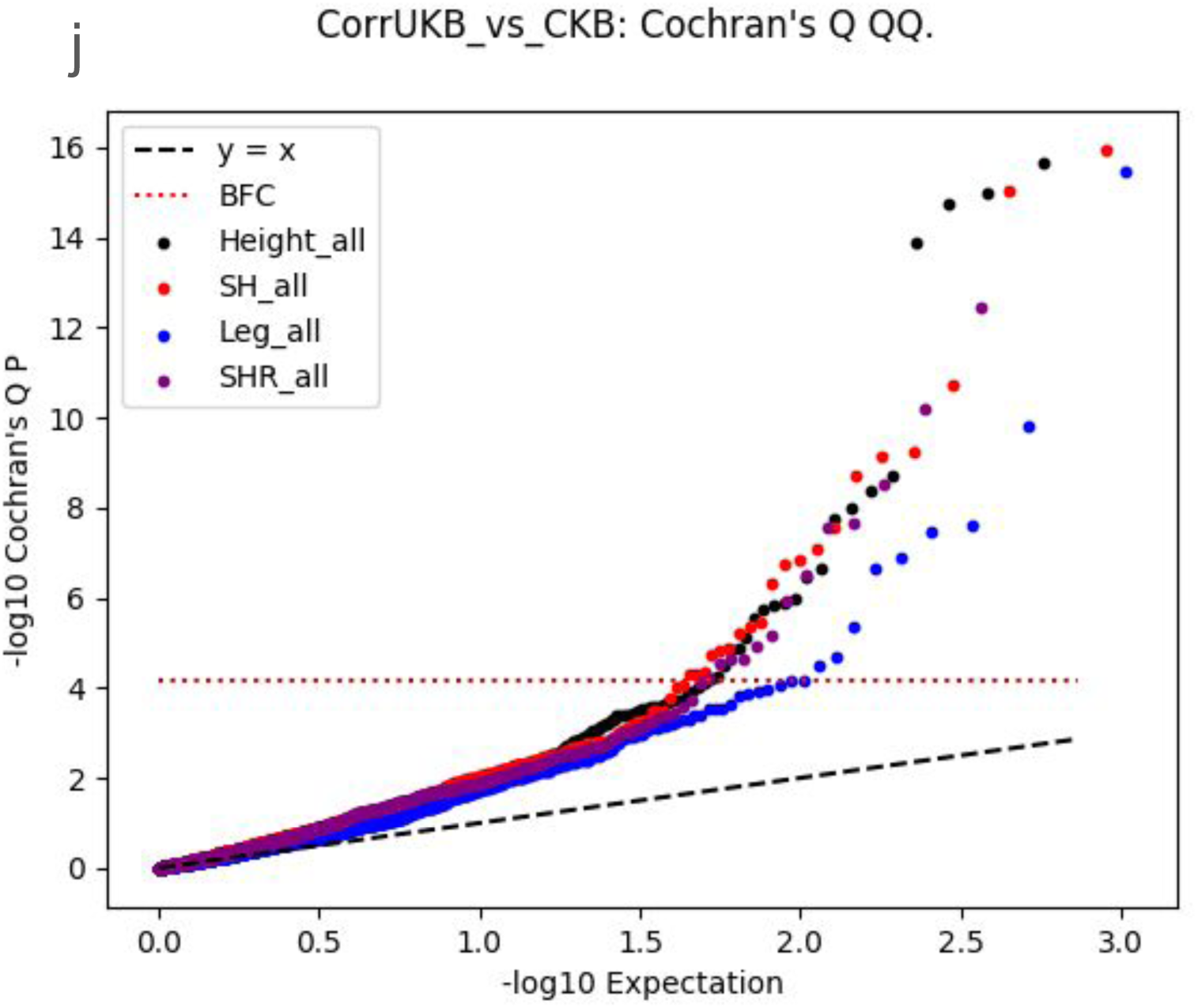
SNPs were discovered in GWAS UKB half 1. A. Effect sizes were corrected for winners curse. SNP effect sizes were on average equivalent when compared to the same SNPs measured in UKB half 2 (red), showing winners curse correction correctly attenuates effect size. SNP effect sizes were on average larger in UKB when compared to the same SNPs measured in CKB (dark blue: UKB half 1 with winners curse correction, light blue: UKB half 2 replication). B. Effect sizes of SNPs discovered in UKB half 1 are larger than when estimating in UKB half 2 (pink), but after winners curse correction, effect sizes are equal on average (red). C. Similar to B, effect sizes of SNPs discovered in UKB half 1 are larger than when estimating in UKB half 2 downsampled to N = CKB unrelated (pink), but after winners curse correction, effect sizes are equal on average (red). D. Similar to B and C, effect sizes of SNPs discovered in UKB half 1 are larger than when estimating in UKB half 2 downsampled to N = CKB unrelated (pink), but after winners curse correction, effect sizes are closer to equal on average (but not equal, red). E. Effect size comparison between SNPs discovered in UKB (uncorrected) and replication CKB; effect sizes are larger in discovery dataset, which is likely due in part to winners curse. F. Similar to E, effect size comparison between SNPs discovered in CKB (uncorrected) and replication UKB; effect sizes are larger in discovery dataset, which is likely due in part to winners curse.

**Supplemental Figure 18:**
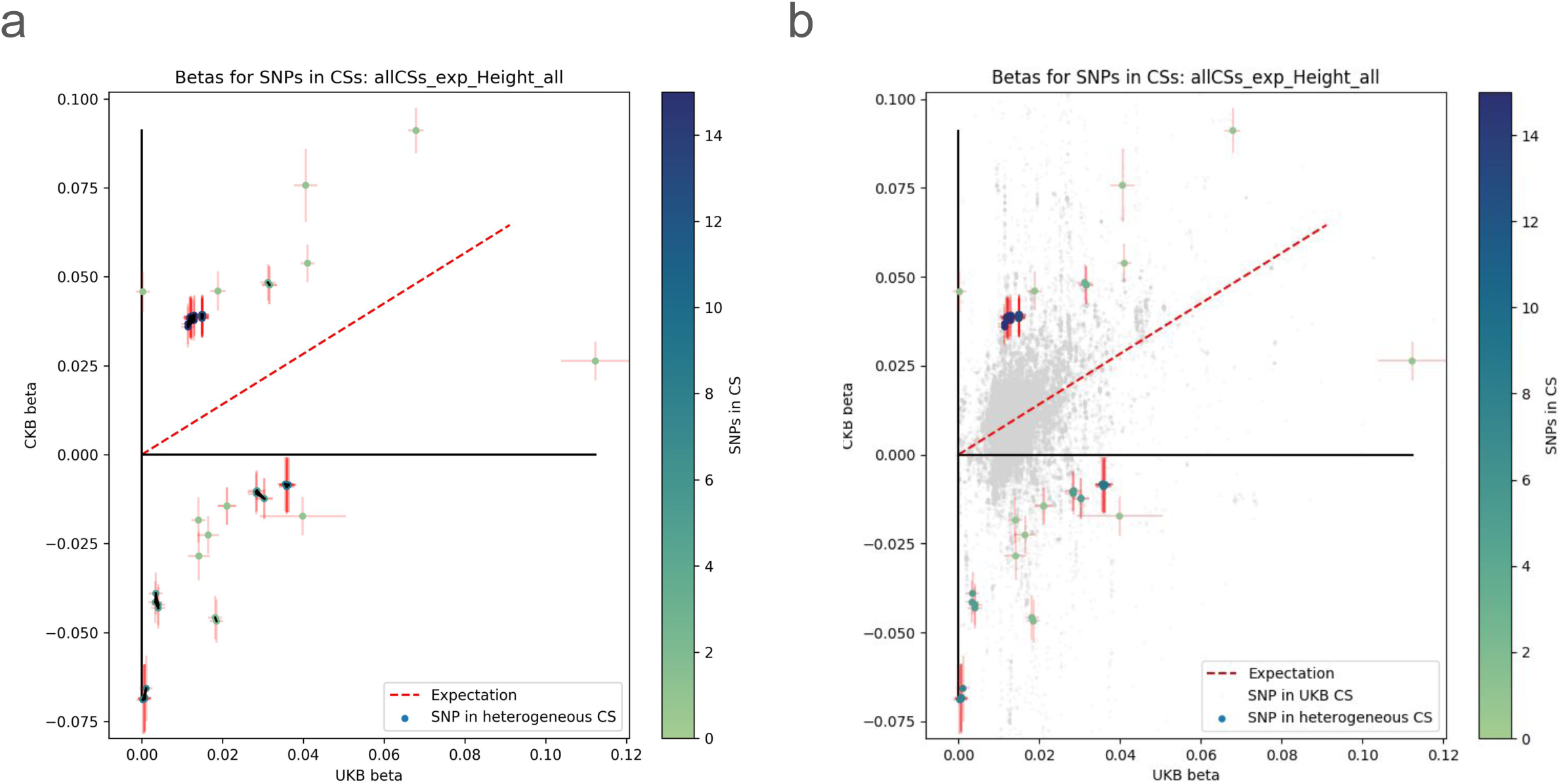
Ancestrally heterogeneous SNP effect sizes [used] A. UKB and CKB effect sizes for only heterogeneous CSs colored by the number of SNPs in each CS. SNPs in the same CS are connected by a black line. B. UKB and CKB effect sizes for all CSs. Heterogeneous CSs are colored by the number of SNPs in each CS.

**Supplemental figure 19:**
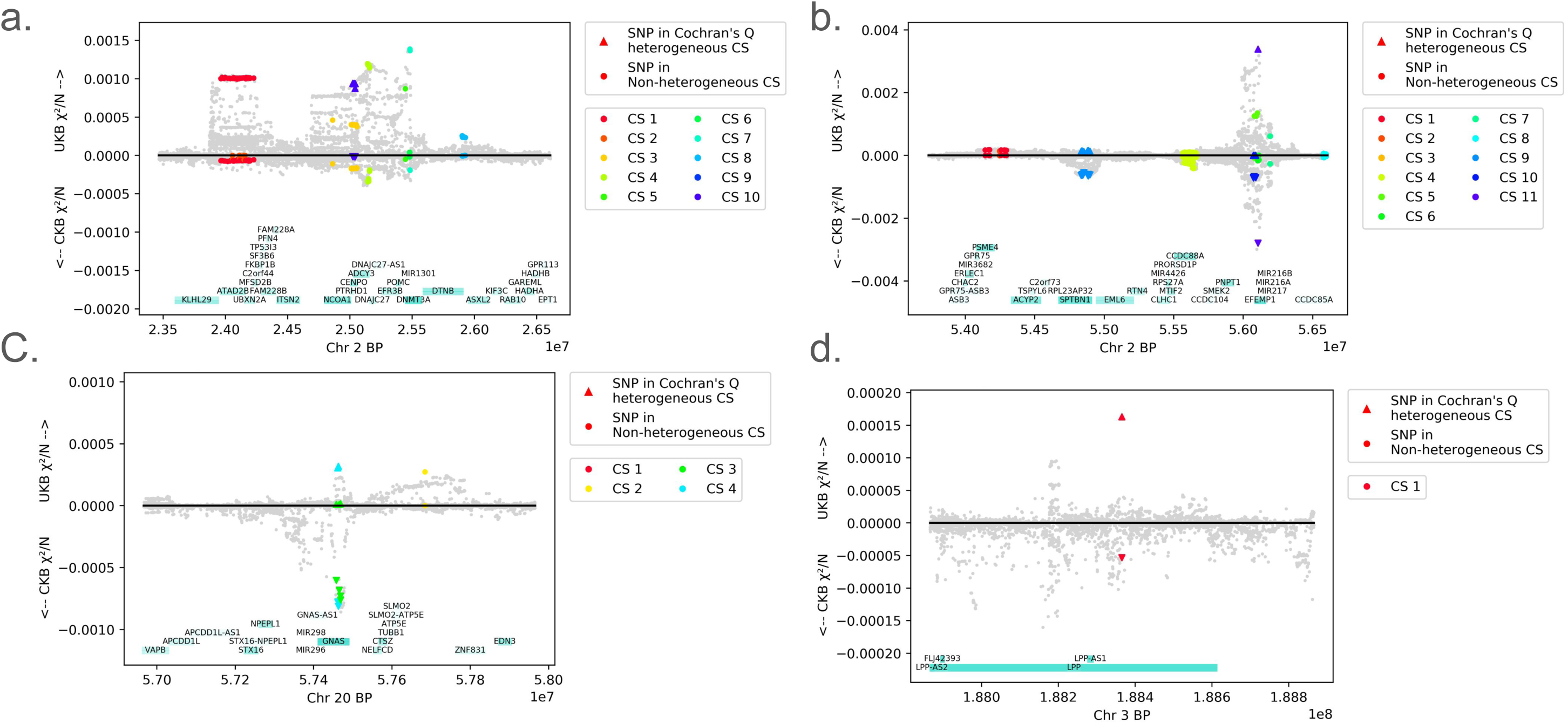
3 loci containing A-het CSs. A. Locus on chromosome 2 ranging 3Mbp contains 10 CSs, 1 of which (CS 10) is A-het. B. Locus on chromosome 2 ranging 3Mbp contains 11 CSs, 3 of which (CSs 8, 10, and 11) are A-het. C. Locus on chromosome 20 ranging 2Mbp contains 4 CSs, 2 of which (CSs 3 and 4) are A-het. D. Locus on chromosome 3 ranging 1Mbp contains 1 CS that is BFC-corrected significant when accounting for only simple loci.

**Supplemental figure 20:**
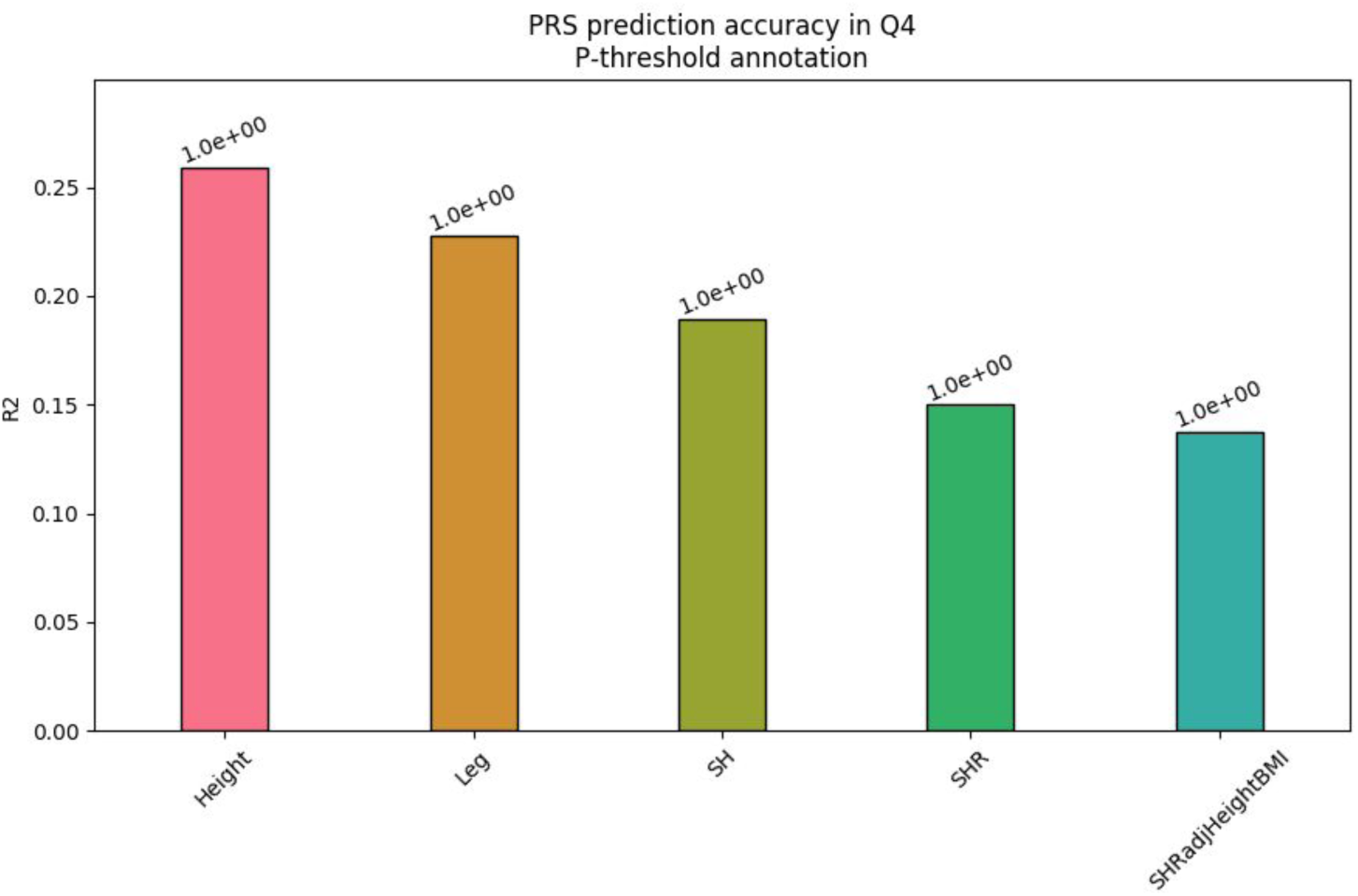
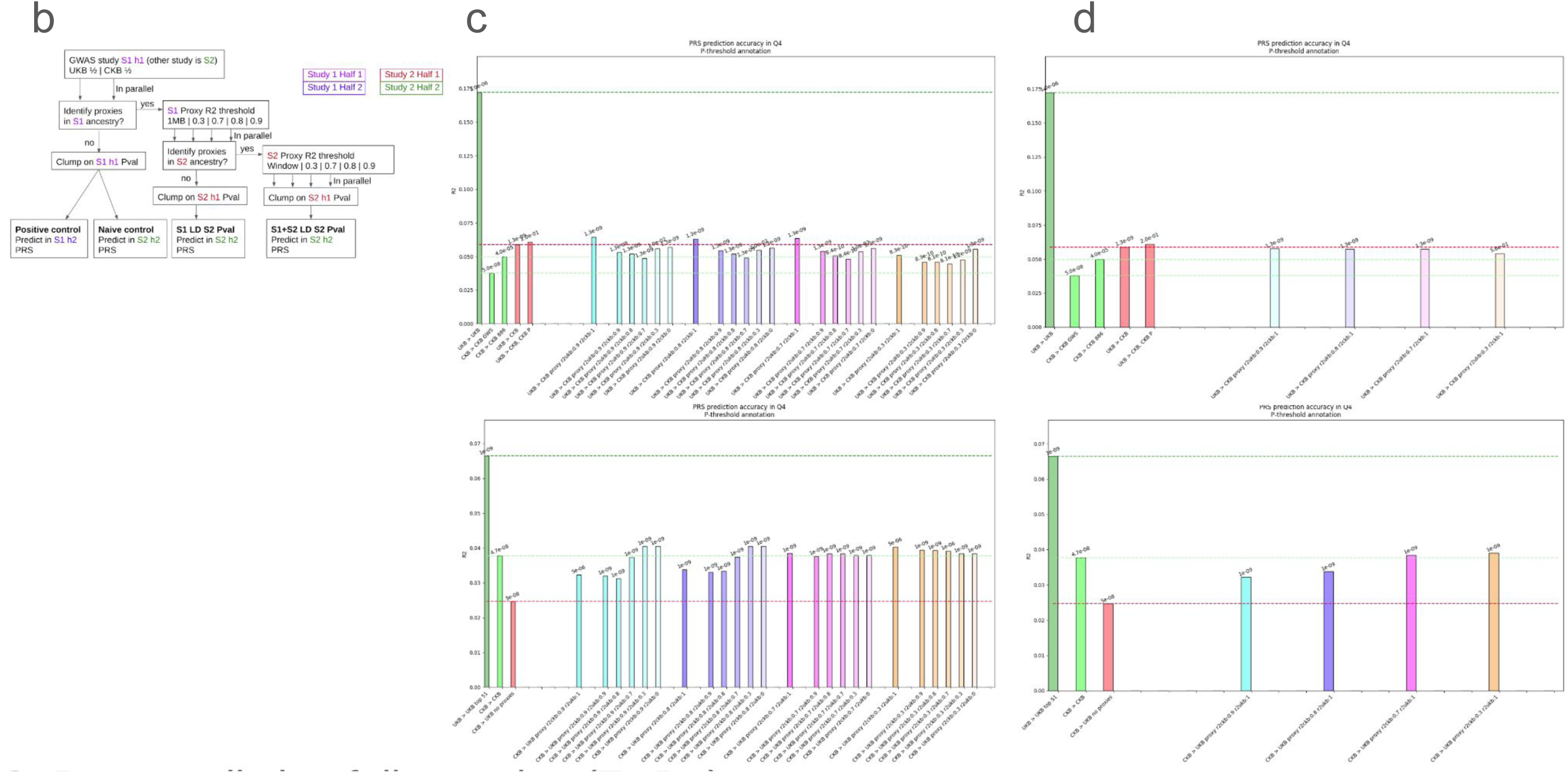
A. PRS using pruning and thresholding performed on body proportion phenotypes in UKB, as measured by prediction R2. PRS for height has the highest R2 tested, and SHR has the lowest (matching behaviour for heritability). Supplemental figure caption: A. Schematic describing approaches for control generation and use of LD and genetic windows to modify SNP selection. B, top. Using UKB GWAS to perform PRS in CKB, measured with prediction R2. For controls, basic pruning and thresholding was applied. For experiments, proxies to lead SNPs were matched at multiple LD thresholds (light blue, dark blue, purple, orange) in both the discovery and the prediction dataset, and then P+T PRS performed. B, bottom. Similar to (top) using CKB GWAS to perform PRS in UKB. C, top. Using UKB GWAS to perform PRS in CKB, measured with prediction R2. For controls, basic pruning and thresholding was applied. For experiments, proxies to lead SNPs were matched at multiple LD thresholds (light blue, dark blue, purple, orange) measured in the discovery dataset, and then P+T PRS performed. B, bottom. Similar to (top) using CKB GWAS to perform PRS in UKB.

